# Phantom Forms in Amblyopic Vision and what they reveal about the Generative Brain

**DOI:** 10.1101/2025.06.24.661078

**Authors:** Akihito Maruya, Bhavatharini Ramakrishnan, Farzaneh Olianezhad, Jingyun Wang, Jose-Manuel Alonso, Qasim Zaidi

## Abstract

What we see is generated by our cortical neurons processing sensory input. Amblyopia provides a perfect case for studying the neural generation of percepts because abnormal neural development of cortex leads to many amblyopes seeing more complex phantom forms than the viewed stimulus through their amblyopic eye (AE) but not through the fellow eye (FE). Using computer guided dichoptic displays, we showed that 92.6% of phantom percepts generated by single gratings shown to AE were matched exactly by plaids (sums of two gratings) shown to FE. A formal equation between the cortical signals generated by the gratings seen through AE and the signals generated by their matched patterns seen through FE, provided the derivation of AE cortical receptive fields as linear transforms of FE receptive fields. The transformed AE cortical model accurately generated the phantoms, was validated via reverse-correlation, and explained deficits in form detection and spatial localization. This study elucidates the close link between percepts and neuronal receptive fields.

**TEASER:** Changes in cortical neurons’ receptive fields can lead to amblyopes seeing phantom features that are not present in viewed images.

## INTRODUCTION

If we could picture the percepts of people (or animals) and generate these images from cortical models (Generative Phenomenology), we could learn so much more than from experimental methods which only provide numerical estimates of parameters. Amblyopia, a disorder of spatial vision, provides an interesting case for a generative study of phenomenology because signals from the two eyes go through partially different cortical neuronal populations. Abnormal neural development in the cortex is a result of imbalance in the input from the two eyes caused by early misalignment (strabismus), lower refractive power (anisometropia), or form deprivation (cataract) of one eye^1^. The disorder persists despite the restoration of good retinal image quality and eye health and affects 2-3% of the population. Consequently, many amblyopes see distorted forms through the amblyopic eye (AE) but not through the fellow eye (FE). AE distortions can potentially be matched perfectly through the FE by rapidly generating images from a large set and using efficient search procedures. The matched images would perform the function of a “Perceptogram”, which is an externally viewable record of an internal perceptual image^2^ or of perceptual distortions of a target image^3^. After obtaining the perceptogram, synthesizing it from generative cortical models can provide detailed insights into cortical processing and neural deficits, and teach us about how forms are created by the brain from the sensory input.

Striking form distortions seen by amblyopes had been noted for letters^4^ and circular shapes^5^, but the most diagnostic distortions were documented by showing high contrast sinusoidal gratings to the AE and drawing the percept through the FE^6^. 25 years after the initial observations, drawings of 20 out of 30 amblyopes were shown to fall into a few gestalts^7^: correct grating orientation with an additional lower contrast grating at some oblique orientation, gratings with abrupt positional shifts orthogonal to the grating orientation, gratings with abrupt positional shifts oblique to the grating orientation, gaps in the grating, fragmented gratings, wavy appearance of straight gratings. The extra features drawn when viewing gratings are phantom percepts and the drawings were shown to resemble plaids formed by sums of pairs of gratings^7^, a key insight reproduced in Figure 1A. Local orientations of shapes and patterns are initially extracted by neurons in primary visual cortex (V1) and then combined in later areas into gestalts. Seeing phantom features is unlike most visual disorders in not being a deficit in perception and requires new types of cortical models that transform the percept by adding non-existent features instead of reducing what is seen. The transformed processing likely starts with orientation encoding in V1^8^ but also involves later mechanisms that decode orientation defined forms^9^. The systematic resemblance of the drawings to plaids ruled out neural scrambling^10,11^ or systematic shifts in the neural map^12-14^ as explanations^7^. Phantom forms generated through one eye are potentially a bigger visual problem than the loss of two lines of visual resolution (the standard definition of amblyopia) for discerning forms and matching inputs from the two eyes for stereo vision, so the complete lack of progress in understanding this problem in the last 22 years is surprising.

**Figure 1.**
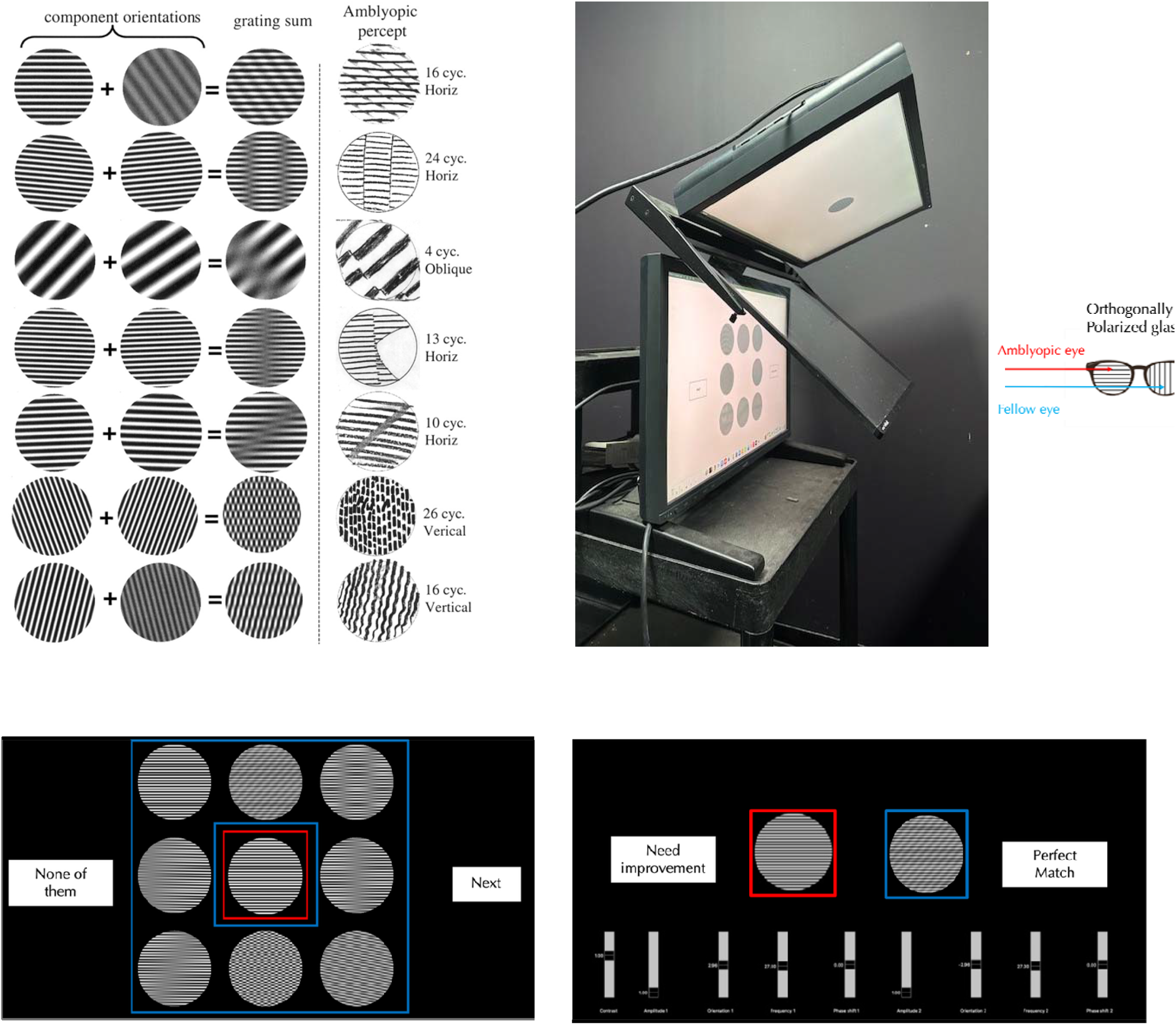
Amblyopic Drawings and Dichoptic Display & Procedure for Perceptograms: **A. Drawings made by amblyopes**^7^: Right: Drawings of seven classes of phantom percepts evoked by the sinusoidal gratings described by # cycles and orientation. Middle: Plaids resembling the drawn phantoms, produced by sums of two component gratings. Left: Component gratings. **B. Dichoptic display using two monitors and a beam splitter (Planar Inc SD2620W):** Images from two LCD monitors with orthogonal polarization are viewed through a beam-splitter that aligns the two images for an observer. The observer wears glasses with the left eyepiece polarized to block one of the images and the right eyepiece polarized orthogonally to block the other. **C. Initial Configuration:** The amblyopic eye (AE) views a central single grating (red border), while the fellow eye (FE) sees a surrounding test grating and seven plaid patterns, each representing a different distortion type (blue borders). The observer selects the surrounding pattern that most closely matches the central grating. **D. Matching Procedure:** The observer fine-tunes four parameters of each of the two gratings within the selected plaid pattern seen by the FE to make a perceptual match to what the AE sees when shown the test grating.

The accuracy of drawings depends on the observer’s skill and the tools provided, and could misrepresent contrast, shading, spatial frequency (SF), orientation (OR), duty cycle, or linearity, so for higher fidelity, we measured perceptograms for 5 amblyopes ages 22-45 using computer generated stimuli. The resemblance of the drawn forms to sinusoidal plaids simplified stimulus generation. Observers viewed a dichoptic display, consisting of two orthogonally polarized LCDs passed through a beam splitter (Figure 1B) and seen through glasses polarized orthogonally so that one screen is seen exclusively by the AE and the other screen exclusively by the FE. The AE was shown a 3 dva (degrees of visual angle) test grating in the center (6, 9 or 12 cyc/deg, 0, 45, 90 or 135° orientations, ON or OFF). Either simultaneously or in alternating sequence, the FE was shown eight 3 dva patterns surrounding the test: 7 plaids matching the different phantom types illustrated in Figure 1A, adjusted to the SF and OR of the test, plus an exact copy of the test grating (Figure 1C). The observer picked the pattern configuration most like the central image, ignoring possible differences in contrast, SF, OR and phase and scotomas. If the observer chose the identical grating, the trial ended. Otherwise, the display showed the AE just the test grating and adjacent to it showed the FE just the initially chosen match (Figure 1D). The chosen plaid was modified to generate better matches by varying contrast, SF, OR and phase of each of the 2 component gratings, until the percept of the test grating seen by the AE was matched perfectly by the generated match in the FE. The procedure was repeated for each grating in three different sessions requiring extensive time from each observer. Since the amblyopic phantoms have been well documented with a large sample^7^, we concentrated on getting complete sets of perceptograms for the three amblyopes in our sample who saw phantoms and deriving a model of amblyopic cortex for each observer that could generate all the images including the veridical percepts.

The 24 matched ON and OFF perceptograms for each observer were used together to infer cortical neural deficits in the ON and OFF systems. We assumed the following linking hypothesis for our model: When the two patterns presented dichoptically look identical to an observer, the signals generated by the test grating seen through AE at some stage of visual cortex match the signals generated by the perceptogram seen through FE. As the simplest instance, we further assumed that AE V1 filters are linear transforms of FE V1 filters. For our model, we separately passed the image of the grating and the perceptogram through orientation-selective Steerable Filters representing a normal V1 cortex and then directly derived linearly transformed filters that collectively generated signals from single gratings that matched signals from perceptograms generated by the unmodified filters. The receptive fields (RF) of the modified filters showed changes in spatial frequency and orientation selectivity but also more complex changes, and a marked increase in spatial extent. The transformed filters can provide a target for physiological measurements, neural development models of amblyopic cortex and for tailoring treatments to ameliorate functional problems caused by amblyopia.

The processing deficits that led to the form phantoms could be adequately explained just in terms of linearly transformed RFs, so we next aimed to explain deficits in other form judgments that would require combinations or correlations of multiple filters. A diagnostic case is judging deviations from a perfectly circular blurred contour when it is replaced with a sinusoidal contour^15^. For a set of fixed frequencies of the distorting waveform, the threshold amplitude for the AE and FE to judge the presence of sinusoidal deviations provides a probe of global form processing^16^. We show that recreating the stimulus from the AE filters derived for our observers, in conjunction with a rotational correlation, predict the change in thresholds for both the AE and FE as a function of modulation frequency, with higher thresholds for the AE, consistent with the published data.

The form phantoms suggest deficits of orientation processing, but there is no direct method to test our models by measuring RFs of single neurons in human visual cortex. A recent reverse correlation study showed broader perceptive orientation tuning for the AE than for eyes of normal observers for vertical gratings^17^, but our perceptograms show that some amblyopes see phantoms and some do not, and neither the phantoms nor the transformed filters are invariant to orientation. Consequently, for all our observers, we estimated perceptive fields^18^ for 0, 45, 90 or 135° orientations of ON and OFF gratings separately for AE and FE using reverse correlation psychophysics^19-20^. Our observers found the task difficult, leading to noisy results especially for the AE, and the procedure cannot identify the cortical locus of the modified filters, but the perceptive orientation tuning through AE was broader, weaker, and shifted compared to FE orientation tuning for the observers who saw phantoms, but not for those who did not, providing additional insight into the orientation deficits underlying the form phantoms.

Note: A major component of this work is mathematical and computational. We explain the operations in words in the main text and present the equations in the **MATERIALS AND METHODS** section in the **Mathematical & Statistical Details** subsection under the same headings as the corresponding text sections.

## RESULTS

### Perceptograms

Three out of our 5 amblyopic observers (Table 1) reported seeing form phantoms in gratings shown to the AE, i.e. 60% versus 67% in the much larger sample^7^. Since each observer’s images take up a lot of space, in the main text we present perceptograms and models from one session each for Observers 3-5 in detail in the main text, excluding the 12 cycles/deg condition for Obs 4, who reported only a uniform gray disk due to the spatial frequency exceeding their resolution limit. Figure 2A shows the 24 gratings and their corresponding perceptograms from a single session. The left four columns display ON gratings, and the right four columns display OFF gratings: the same gratings are ON or OFF depending on being increments or decrements from a black or white background respectively. Each orientation-frequency combination produced a distinct phantom pattern. Stimulus polarity also influenced perception: for example, at 90° and 6 cycles/deg, for Observer 3, the ON grating produced a net-like appearance, while the OFF grating yielded a smoother, wavy structure with shadows at the top and bottom, suggesting that deficits could be different for the ON and OFF systems.

**Table 1.** Observer details.

| Subject | Age | Gender | Type of amblyopia | Visual Acuity (LogMAR) | Refraction | Fixation | Distortio |
| --- | --- | --- | --- | --- | --- | --- | --- |
| Obs 1 | 24 | F | Anisometropic | OD: 0.00<br>OS: 0.20 | OD: +6.75/-0.25 X 125<br>OS: +7.75/-1.00 X 075 | Central | No |
| Obs 2 | 49 | F | Combined | OD: 0.20<br>OS: 0.00 | OD: +5.50<br>OS: +5.00 | Central | No |
| Obs 3 | 22 | F | Anisometropic | OD: 0.00<br>OS: 0.30 | OD: -2.50/-1.00 X 180<br>OS: +0.75/-1.00 X 180 | Central | Yes |
| Obs 4 | 22 | F | Combined | OD: 1.00<br>OS: 0.00 | OD: -22.00/-2.00 X 035<br>OS: -11.50/-2.75 X 140 | Temporal | Yes |
| Obs 5 | 23 | F | Anisometropic | OD: 0.40<br>OS: -0.10 | OD: 0.00/-0.50 X 018<br>OS: -3.25/-0.50 X 121 | Central | Yes |

**Figure 2.**
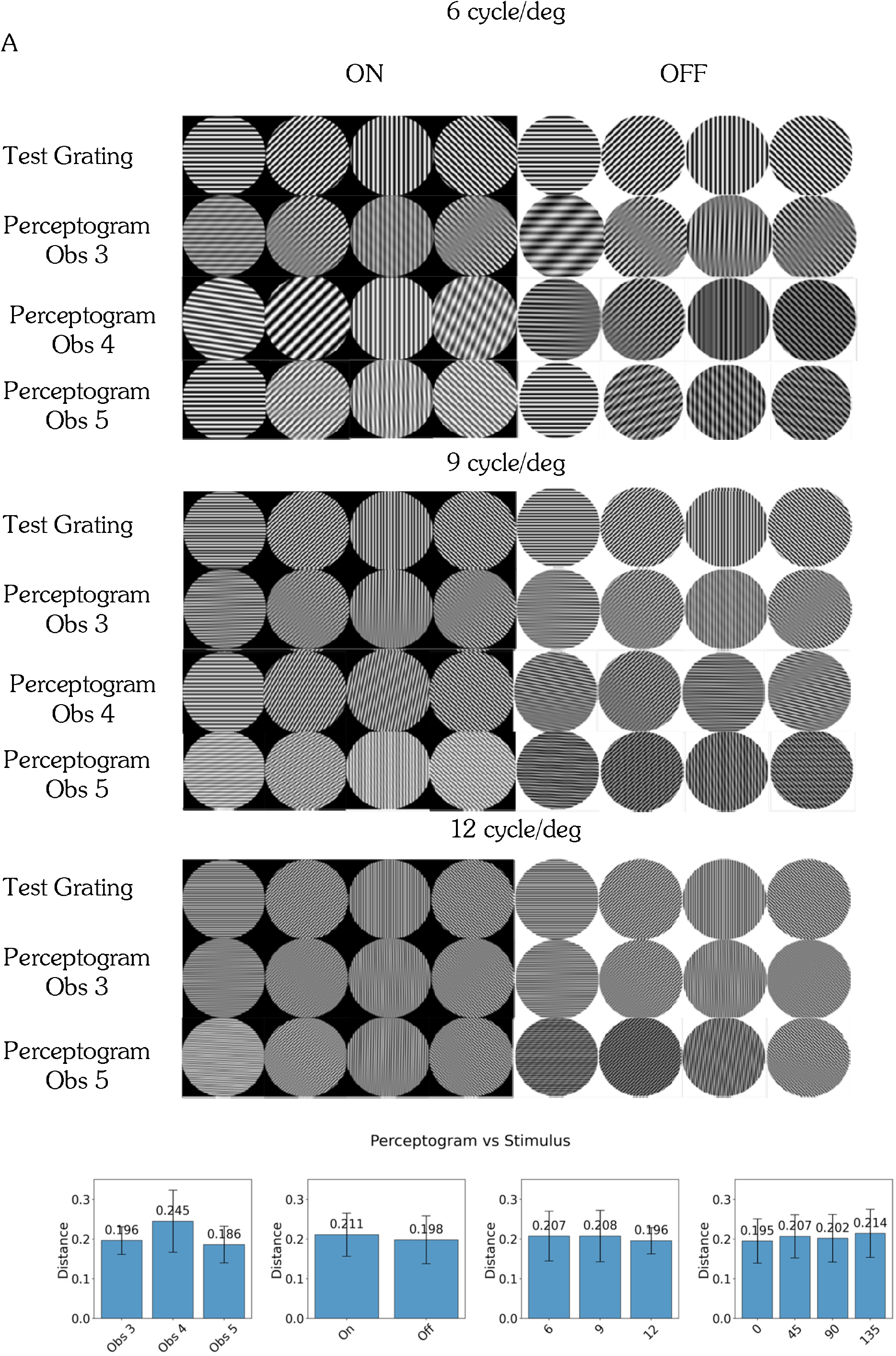
Perceptogram Images: A. Eight orientations of ON and OFF gratings per spatial frequency (6, 9, 12 cyc/deg) shown to AE and corresponding perceptograms obtained by matches in FE are shown for Obs 3, 4, 5 for one session each. Obs 4 could not resolve 12 cyc/deg. B. DISTS estimated perceptual distance between measured perceptograms and stimulus gratings.

Complete sets of perceptograms from all three observers who saw phantoms are shown in Supplemental Figure S1. Across observers, the most frequently reported percepts resembled a grating with one or more blurred segments separating phase-shifts, like those illustrated in rows 2–5 of Figure A1. These patterns emerged from the combination of two gratings with closely matched spatial frequencies and contrast, but with slight differences in orientation and phase. The second most common patterns were net-like, resulting from the combination of one grating like the test and another with a lower spatial frequency, reduced contrast, and a different orientation (top row Figure A1). The bottom two patterns arise when the two constituent gratings are quite different in orientation from the test grating, with an interrupted pattern if they are the same contrast, and a wavy pattern if one is of lower contrast. The estimated perceptual distance between the perceptograms and the test gratings averaged over all the conditions for the three observers who saw phantoms was quantified as 0.204 and significantly different from 0.0 that denotes a perfect perceptual match by the Deep Image Structure and Texture Similarity (DISTS) metric^21^. The average perceptual distances between observers, spatial frequencies, orientations and ON/OFF polarities were not significantly different (Figure 2B).

The summation of two sinusoidal gratings failed to fully capture the phantom percept only in 16 out of the 216 cases (7.4%). In these cases, observers verbally reported the mismatches—generally describing contrast mismatches. When contrast mismatches were insufficient descriptions, observers provided additional verbal descriptions (included in the far-right column of Figure S1), often reporting irregularities across the patterns. When verbal descriptions were inadequate, observers illustrated their percepts using Procreate on an iPad, after being trained to control linearity, width, contrast, shading and orientation to make these drawings much more representative of the percept than the previously published pencil drawings. A few drawings showed greater waviness than captured by the plaids such as in the bottom row of Figure 1A. A careful drawing could take more than 30 minutes, making it unfeasible to use that method for the complete set of stimuli.

Across sessions, perceptograms were largely consistent within observers, with most variations reflecting minor differences in phase alignment. However, occasional session-to-session variability in reported pattern type was observed. For example, for ON gratings oriented at 0° with 6 cycles/deg frequency, Obs 3 reported a net-like pattern in two sessions but described a grating with a shadow in the third. The exact cause of this variability remains to be investigated.

The 92.6% of phantoms that were accepted as matched by the digitally generated plaids, along with the verbal descriptions and Procreate drawings of the rest, demonstrate the importance of measuring the perceptograms, because it seems that the lack of gradual shading in the published drawings leading to depictions as lines or square wave gratings, sometimes with unequal duty cycles, are due to limitations of drawing skills with pencil and paper and do not represent perceived phantoms. In addition, unlike the incomplete set of published drawings, we have complete sets of perceptograms for each observer, allowing us to build models that simultaneously account for both form phantoms and veridical percepts. Therefore, we build cortical models that reproduce perceptograms instead of the drawings. If the irregularities and waviness were frequent, we would have incorporated neural scrambling and under-sampling in the models, but the occurrences were too rare.

### Cortical model

We began with the working hypothesis that the form phantoms observed in amblyopia result from receptive field (RF) alterations in cells of early visual cortex that get input from the AE while the RFs are normal for cells that get input from the FE, with later visual areas processing the signals similarly irrespective of eye of origin. To model the responses of the FE driven normal V1 cortex to the stimuli, we used steerable pyramids for frequency decomposition of input images in the Fourier domain^22^. This method is translation- and rotation-invariant, prevents aliasing, and allows for perfect reconstruction of the original image. The Supplemental file gives the mathematical details. The input image is first decomposed into high-pass and low-pass components by multiplying its Fourier transform with a high-pass filter that emphasizes higher spatial frequencies, and a low-pass filter that retains lower frequencies. Both filters are selective only for spatial frequency and are independent of orientation. The low-pass band is subsequently decomposed into oriented bandpass filters. The process is repeated across the specified number of scales. To visualize the spatial profile of these filters, a centered delta function was passed through each filter and the inverse Fourier transform applied. The V1 filters for the FE are shown in Figure 3A for 16 orientations and four spatial scales arranged from high frequency (top) to low (bottom). The resulting patterns resemble oriented Gabor filters in being soft windowed versions of sine functions, and by passing perceptograms through the complete set of filters, we ascertained that they satisfy perfect reconstruction.

**Figure 3.**
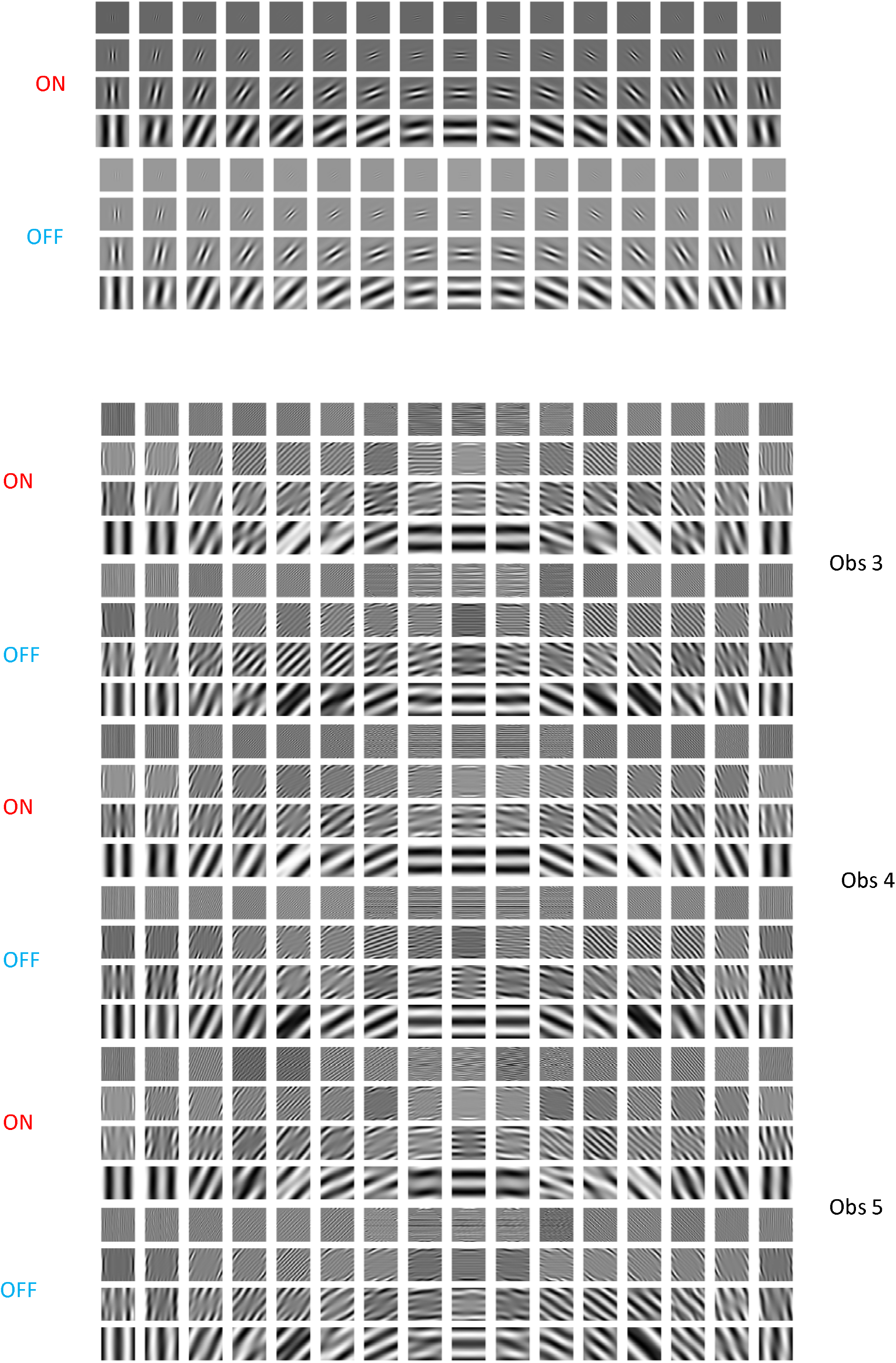
Cortical Model for Amblyopia. **A: Fellow Eye Filters:** Visualization of 4 orientated sub-bands from a standard steerable pyramid, representing the FE cortex, showing the response to a central impulse for ON and OFF filters that are identical except for opposite light-dark polarity. **B: Amblyopic Eye Filters:** Visualization of each orientation sub-band from the linearly transformed amblyopic steerable pyramid in response to a central impulse, representing the AE ON and OFF cortical RFs for Obs 3, 4 & 5.

Our linking hypothesis is that the cortical representation of the test grating through AE neurons matches the cortical representation of the perceptogram through FE neurons. We tested if linear transforms applied to all standard oriented band-pass filters can collectively account for the cortical transformations, which allows us to write a formal equation equating the signals from the gratings through the linearly transformed filters to the signals from the perceptograms through the normal filters (Supplemental file: Cortical model). The supplement shows that this equation can be simplified to directly derive the linear transformation from the ratio of the cross-spectrum to the power-spectrum. As examples, the transformed AE filters for Obs 3-5 saw phantom forms for one session each for the ON and OFF conditions are visualized in Figure 3B. Comparing the filters in Figures 3A and 3B shows that some of the AE RF distortions could be changes in spatial frequency and orientation tuning of V1 cells, corresponding to linear operations of scaling and rotation, but others seem to create complex non-Gabor RFs suggesting combined operations including shearing, particularly for diagonally oriented components. The transformed filters also exhibit an increase in effective spatial extent as the ON and OFF lobes repeat over the complete 3 dva of the stimulus. The model cannot specify if these changes are in V1 RFs or in later combinations of V1 outputs. In particular, the greater spatial extent and some of the more complex changes could reflect broader spatial pooling of V1 outputs.

We built models that covered all three spatial frequencies and four orientations, but separately for each observer, session, and stimulus type (ON vs. OFF). To test if each discrete set of linearly transformed filters is adequate, we applied the complete set in the Fourier domain (including the untransformed and unoriented high-pass and low-pass filters) to the Fourier transforms of the grating set, and then took the inverse Fourier transform to generate the corresponding perceptograms. Example comparisons are shown in Figure 4A for the Obs 3 perceptograms from Figure 2A. Within each pair, the top image depicts the observer’s perceptogram, while the bottom shows the corresponding reconstruction from the AE model. Comparisons for all perceptograms for all observers are presented in Supplemental Figure S1. Visual inspection shows that each model accurately predicts the observer’s percepts with remarkable fidelity. In addition, the DISTS^21^ estimated perceptual distance between measured perceptograms and their model generated recreations averaged over all the conditions for the three observers who saw phantoms was 0.004 and not significantly different from 0.0 that denotes a perfect perceptual match. The average perceptual distances between observers, spatial frequencies, orientations and ON/OFF polarities were all not significantly different from 0.0 (Figure 4B).

**Figure 4.**
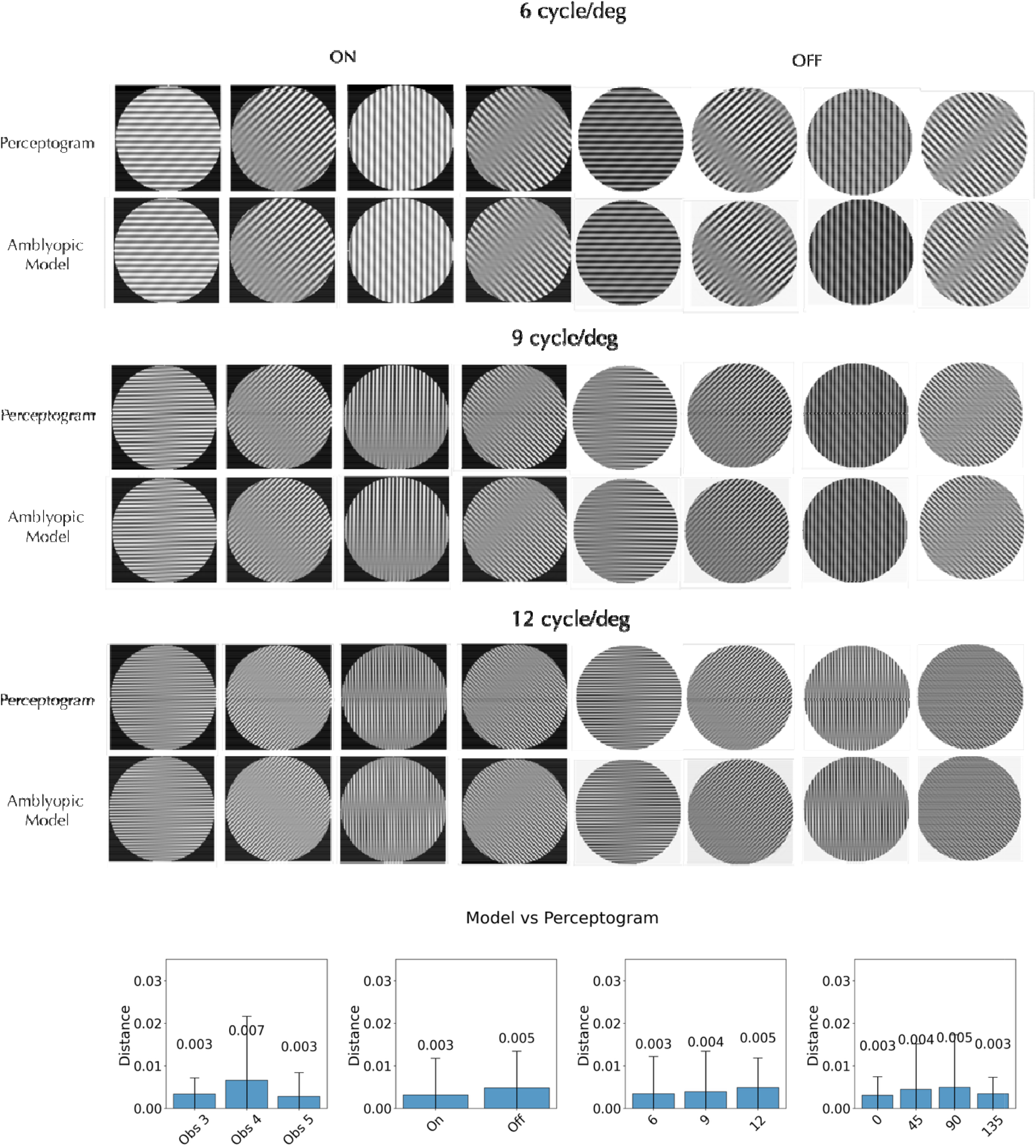
Model reconstructions of Perceptograms and comparisons of estimated perceptual distances between images: A. Perceptograms for eight orientations of ON and OFF gratings per spatial frequency (6, 9, 12 cyc/deg) are shown for Obs 3 for one session along with the corresponding reconstructions from the linearly transformed V1 filters representing the AE cortex. Recreations for perceptograms for all three observers from each of their 3 sessions are shown in the Supplement. B. DISTS estimated perceptual distance between measured perceptograms and their model recreations. The scale is 10 times finer than in Figure 2B. The model accurately generated the perceptograms.

### Sinusoidal Deformations from Circularity

We tested our model on results of a task claimed to have a global component: the detection of sinusoidal deviations from circularity of the fourth derivative of a Gaussian (D4)^16^. The left panel in Figure 5A shows the circular D4 stimulus and its log power spectrum, while the right panel shows one of four sinusoidally deformed D4s, each with a radial frequency of 4, 6, 8 or 10 cycles per 360° with its corresponding log power spectrum. The circular D4 exhibits uniform power across all orientations, whereas the sinusoidally modulated D4s reveal localized increases in power at specific orientations, producing the same number of ridge-like structures in the spectrum as the number of cycles per 360°. As modulation amplitude increases, the local differences from the circular D4 (far left) become more perceptually apparent, and the ridges in the power spectrum more pronounced. Detecting these orientation-specific enhancements is critical for identifying deformation, but there is also a curvature detection aspect to this task beyond judging local orientation deviations. When the circular and modulated D4 stimuli were presented simultaneously, and the detection threshold for the modulation was measured separately for the AE and FE^15^, detection thresholds for FE decreased with increasing radial frequency, but the AE thresholds were fairly constant and consistently higher than FE across all tested frequencies.

**Figure 5.**
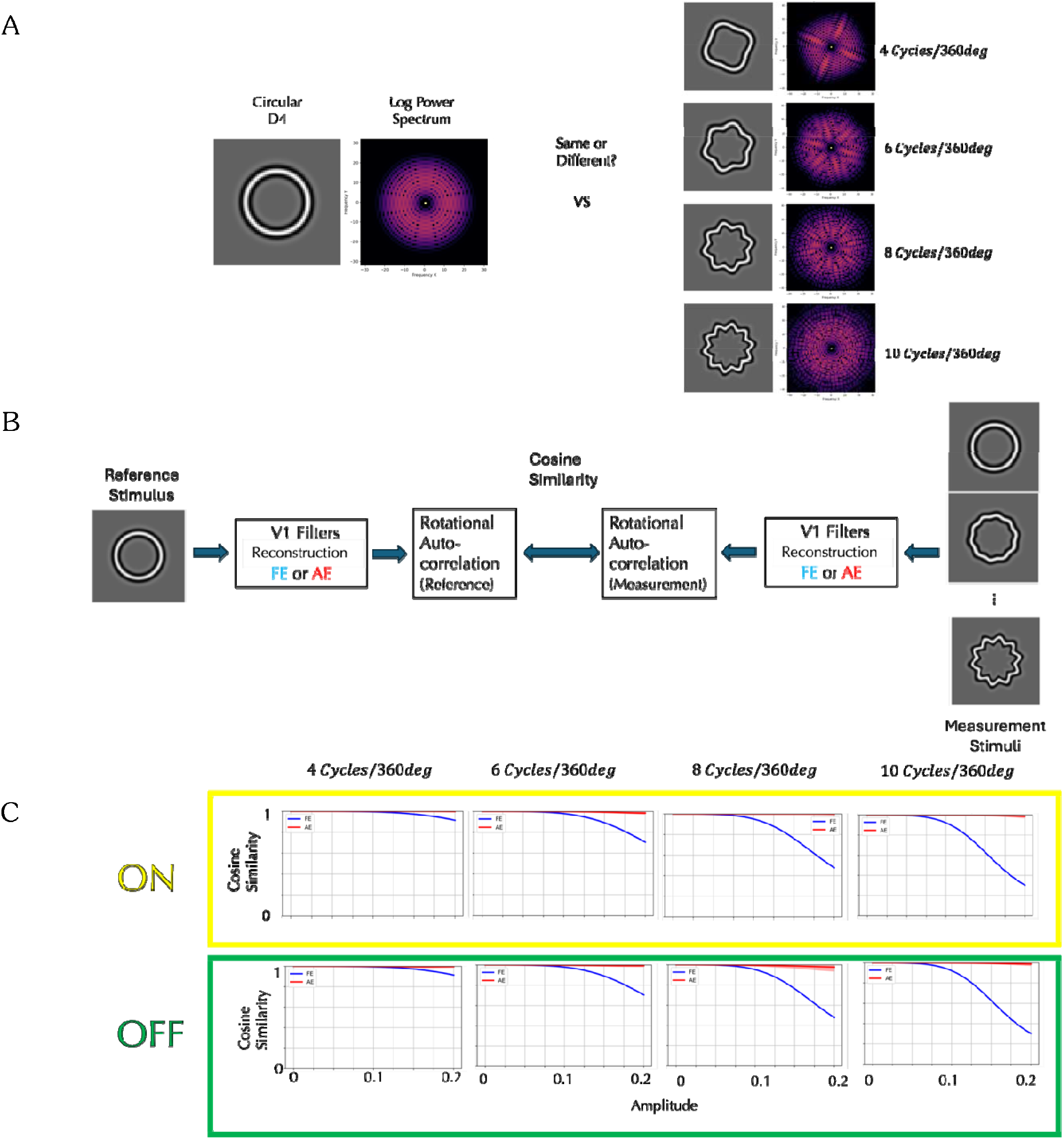
Detection of Sinusoidal Deformations from Circular D4 Stimuli and Model Simulations: **A:** Circular and deformed D4 stimuli and their log power spectra. While the circular D4 displays uniform power across orientations, the deformed D4s show localized increases in power at specific orientations, forming ridge-like structures. Amblyopic eyes consistently exhibited higher amplitude thresholds across radial frequencies than fellow eyes. **B:** Model for responses of the fellow eye and amblyopic eye. The statistical representation of the circular D4, processed through either the fellow eye (FE) or amblyopic eye (AE) V1 filters and then through a rotational correlation, is compared to those of the modulated D4 stimuli at various amplitudes using cosine similarity. **C:** Yellow box shows predictions from the ON amblyopic model, and Green box from the OFF amblyopic model, averaged over 3 observers who saw distortions (shading indicating ±1 standard deviation). Each panel plots cosine similarity as a function of amplitude for one radial frequency (4–10 cycles/360°), where a higher similarity leads to a higher detection threshold. For the FE (blue curves), cosine similarity decreases with increasing amplitude, and this reduction is more pronounced at higher radial frequencies. The AE curves (red) show relatively high cosine similarity across amplitude and radial frequencies. Both are consistent with published detection thresholds.

To investigate whether our AE V1 model can account for the observed difference in detection thresholds, we constructed a computational model with the schematic architecture shown in Figure 5B for the FE and AE simulations. In the fellow eye (FE) model, the circular D4 was processed through a simulated normal V1 pathway, and then we computed rotational correlations of the reconstructed image derived from the filter responses for rotation angles from −17° to 17° in 1° increments, yielding a feature vector of length 35. Feature vectors were computed similarly for modulated D4 stimuli with varying amplitudes and compared to the reference vector using a cosine similarity metric. In the amblyopic eye (AE) model, the same procedures were applied, but the stimuli were processed through the AE cortical model. This approach allowed us to quantitatively assess how closely the modulated stimuli at each amplitude resembled the circular reference under each cortical model.

The model predictions are presented in Figure 5C. Each panel represents a different radial frequency and compares the ON and OFF AE models to the FE models. Within each panel, cosine similarity is plotted as a function of modulation amplitude. Higher cosine similarity indicates greater representational overlap between the modulated and reference stimuli, which predicts a higher detection threshold. In the FE cortex model, cosine similarity systematically decreases as the amplitude of modulation increases, and the rate of decrease is greater at higher radial frequencies, reflecting that cortical responses show increased sensitivity to higher-frequency deformations. These findings are consistent with the psychophysical results which demonstrated lower detection thresholds for higher-frequency deformations. In contrast, the AE cortex model exhibits relatively high cosine similarity between the modulated and reference D4 stimuli for all amplitudes and radial frequencies. This pattern is consistent with consistent with the higher and constant thresholds reported for AE and explains the diminished sensitivity to form deviations in the amblyopic cortex. This insensitivity is generated by both ON and OFF models of the AE cortex, although the OFF model tends to produce slightly lower cosine similarity values than the ON model, predicting a modest difference in form perception between the two pathways.

### Orientation Tuning via Reverse Correlation

The AE filters for amblyopes who see form phantoms show complex differences from the FE filters, but the perceptograms and D4 results clearly point to deficits in orientation processing. To understand differences in orientation tuning between amblyopes who see phantoms and amblyopes who do not, we measured perceptive fields using reverse correlation (RC)^18-20^ for target gratings of 6 cyc/deg with 0°, 45°, 90°, or 135° orientation for ON and OFF conditions, separately for AE and FE (see Methods: Reverse Correlation). Participants fixated on the central display as a series of orientations were flashed and were instructed to press a button as soon as they detected the target orientation that had been shown to the FE. Given the difficulty that our amblyopic observers had doing the task, we used presentations of 100 msec, which is slower than usual. The Supplemental file describes in detail how we obtained the orientation tuning curves shown in Figure 6. Figure 6A (solid outlines; yellow for ON, green for OFF) depict responses from the two amblyopic observers who did not report form phantoms. Figure 6B (dashed outlines) shows tuning curves from the three amblyopic observers who reported perceiving phantoms. Each row corresponds to a different target orientation, and within each panel, response magnitude is plotted against the physical orientation of the stimulus. Red and blue dots represent the mean responses (based on 5,000 bootstrap resamples) for the amblyopic and fellow eyes, respectively. Solid curves denote the best fitting von Mises functions, with shaded bands indicating ±1 standard deviation. Amblyopic and fellow eye tuning curves are roughly similar for the two amblyopic observers who did not report form phantoms, but the tuning curves from the three amblyopic observers who reported perceiving phantoms clearly show that the AE has reduced peak responses at 0° and 90°, broader tuning at 90°, and marked peak shifts at 45° and 135°. There are some differences between ON and OFF results as the ON curves are slightly broader, but they do not provide sufficient evidence that either system is more affected.

**Figure 6.**
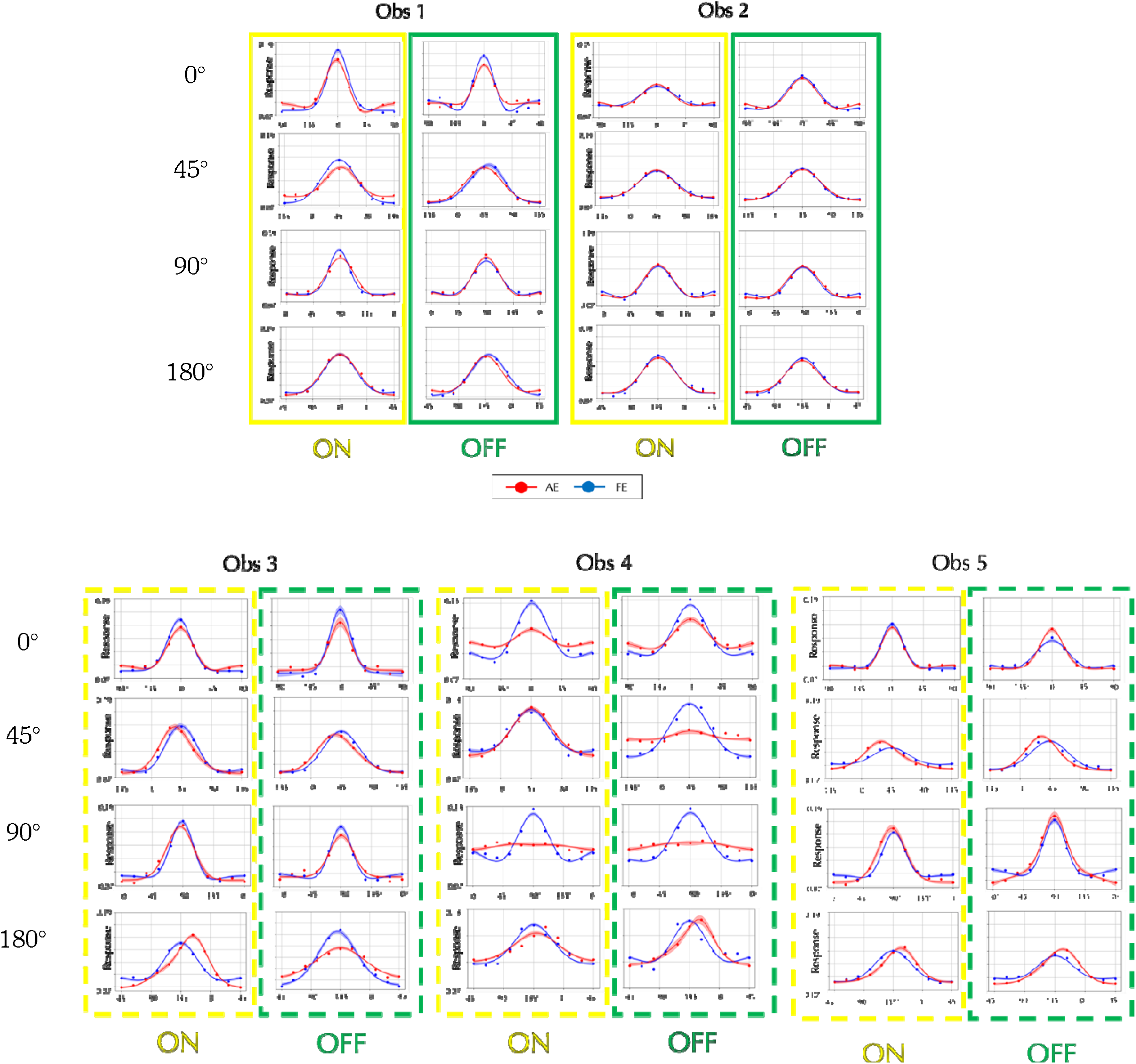
Perceptual Orientation Tuning Curves from Reverse-Correlation for all Observers: A. Results for two amblyopes who did **not** report form phantoms (Solid boxes Yellow: ON, Green: OFF). B. Results for three amblyopes who **did** report phantoms (Dashed boxes). Each row corresponds to a different target orientation. Within each panel, response magnitude is plotted against physical orientation. Response magnitude is scaled to give unit area under the curve. Red and blue dots represent the mean responses (after 5,000 bootstrap resamples) for the amblyopic and fellow eye conditions, respectively. Solid traces are the best fitting von Mises functions, with shaded areas indicating ±1 standard deviation. Most of the mean R-squared values were 0.98 or higher.

## DISCUSSION

This paper solves a 50-year-old puzzle which has great significance for understanding amblyopic vision, and designing potential treatments, but the most significant conclusion comes from the generative part of the model showing that changes in neuronal receptive fields have profound effects on perception, to the extent that observers can see more features than are present in the viewed stimulus. The phantoms/distortions persist despite amblyopes having years to potentially adapt to the transformed RFs. The obvious inference is that receptive fields in normal visual cortex are exquisitely tuned for perception to be veridical in most cases.

The efficiency of orientation processing has been linked to the anisotropy in the distribution of image orientations in natural scenes being matched by the distribution of orientation tuned cells in V1^23^. The less-than-optimal consequence is that the anisotropy in the distribution of orientation and direction tuned cells, leads to anisotropy in the estimates of angles^9^, orientation flows^9^ and optic flows, which makes high level percepts of 3D shapes^9^ and object non-rigidity^24^ orientation-dependent, but leaves forms recognizable. Distortions of receptive fields seem to be more disruptive of perception than variations in distributions of neuronal properties.

Amblyopia is clinically identified by a reduction in the visual acuity of the AE and disrupted binocular function, and these have been the focus of most research, but deficiencies in several low- and high-level perceptual abilities have also been documented. The phantoms, on the other hand, cannot be classified as reduced vision. We definitively confirmed the published perceptual phantoms with perceptograms and showed that they can be explained by linear transformations of V1 RFs. Many transformed RFs resembled V1 RFs albeit with broader tuning and shifted peak^25^, but others were much more complex and may reflect combinations of V1 outputs at later stages, compatible with orientation preference heterogeneity in Amblyopic V2^26^. The transformed RFs can be used as predictions to be tested electrophysiologically for V2 neurons.

Many of the linearly transformed filters in Figure 3B have many more lobes than the normal filters, which gives greater summation area for each RF. This RF structure implies that the gratings that fill the whole RF while matching spatial frequency selectivity would evoke the maximum responses, unlike for normal RFs, which could partially explain the much larger increase in contrast sensitivity for amblyopes as compared to normals as field sizes are increased from 0.25 dva to 1.0 dva^27^, although the greater increase up to field sizes of 8 dva would probably be due to enhanced lateral connections. Extra lobes also shrink the spatial frequency bandwidth and hamper the localization of spatial position^48-50^. The neural development of such abnormal RFs requires a larger spatial extent of inputs, whether it is thalamic inputs to V1 or V1 inputs to V2. Visual blur effectively reduces stimulus contrast, which would increase both the spatial footprint of lateral connectivity and its effect on visual responses ^28^. During the amblyopic critical period, blur has been shown to reduce neuronal contrast sensitivity^29^, but could also spread the formation of connections, leading to extended lateral connections, which could be tested physiologically as part of the development of transformed RFs.

The formal equation between the AE response to gratings and the FE/V1 response to their matched perceptograms, enabled us to derive the transformed AE filters, but does not address their cortical loci. In principle the equation could be used to linearly transform a set of normal RFs in any cortical area. If V2 RFs were as well established as V1, we could have also applied the equation to V2, especially since amblyopia has been shown to disrupt RF structure in V2 neurons^25,26^ possibly even more than it does in V1^27^. However, while it is known that V2 RFs are weighted combinations of large numbers of orientation tuned V1 RFs, and thus have much more intricate internal structure^30^, which corresponds to heightened sensitivity to combinations of discontinuous orientations^31^ and textures^32^, there is no established canonical form that we could have used in the model.

The transformed AE RFs were not sufficient to explain the reduced sensitivity to the global judgement involving sinusoidal perturbations of contours, but required the addition of rotational correlation, which is more general than a model involving curvature specific neuronal connections^33^. Computational models of V2 neuronal sensitivities to naturalistic textures include many different correlations between V1 outputs^23,32^, and it is possible that some V4 neurons that are sensitive to concentric stimuli^34^ incorporate rotational correlations between earlier outputs.

The perceptual phantoms in the perceptograms consist of multiple oriented gratings which suggest broader tuning of orientation selectivity. Consistent with this idea, the results of the reverse correlation experiment showed orientation tuning differences between AE and FE for amblyopes who saw phantoms, but not for those that did not.

Amblyopia is generally thought to result from relative monocular deprivation of one eye causing a shift in ocular dominance of binocular neurons in V1 during an early critical period^35,36)^. However, multiple instances of abnormalities in areas beyond V1^28,29^ require more than a shift in ocular dominance^37^. Our results suggest that other mechanisms of neural development of distortions in RFs caused by amblyopia need to be investigated. Our initial conjecture on starting this project was that the development of frequency and orientation selective V1 RFs requires balancing ON and OFF thalamic inputs^38^, so if they are not balanced, RFs will not develop normally. We had also found some evidence that the ON system is affected more than the OFF in amblyopia, as measured by grating resolution and salience of whites versus blacks in random checkerboards^39^, buttressing our idea. That provided the motivation to use separate ON and OFF gratings. The perceptograms were sometimes different for the two polarities for gratings of the same orientation and frequency, but they showed only a slightly stronger effect of amblyopia for ON versus OFF stimuli, and the effects were mixed in the reverse correlation results. In the perceptogram measurements, the stimuli were seen for prolonged periods with eye movements and may not have isolated ON and OFF systems very well, but the reverse correlation stimuli were flashed briefly, so they truly are increments and decrements. In addition, a comparison of the AE filters to FE filters in Figure 2 reveals that whereas some RFs do become less selective in the AE, the transformations in other RFs are too complex to be due solely to a weaker ON or OFF system. Our results thus do not provide strong support for our initial conjecture, and the neural development of abnormal RFs remains an open question, with broader connectivity caused by reduced contrast signals during the critical period being an important candidate worth testing.

It is worth considering the functional implications of our results. Stereo is compromised in amblyopia, so amblyopes have to rely on monocular cues for perception of 3D scenes and objects. However, many monocular cues are critically based on orientation information, so if oriented patterns are perceived as distorted the brain may not be able to compensate. There is strong evidence for orientation information being critical for 3D shape from texture^40-42^, mirror symmetry^43,44^ and 3D pose perception^45^. Figure 7 shows that whereas the diagnostic orientation flows in a 3D shape from texture stimulus^41^ are retained by the FE cortical model, they are lost by the AE model. Simulations using mirror symmetric complex images^43^ showed loss of detail by the AE model and lack of symmetry in the output for some but not all images. Our models predict that the amblyopic eye will show greater deficiencies in these functions. It is possible that if the AE models were derived from a more diverse sample of perceptograms, the filters may extract more information from such images but investigating performance on these tasks for amblyopic observers that do and do not see form phantoms would be worthwhile, as it could also test whether the non-amblyopic eye is sufficient.

**Figure 7.**
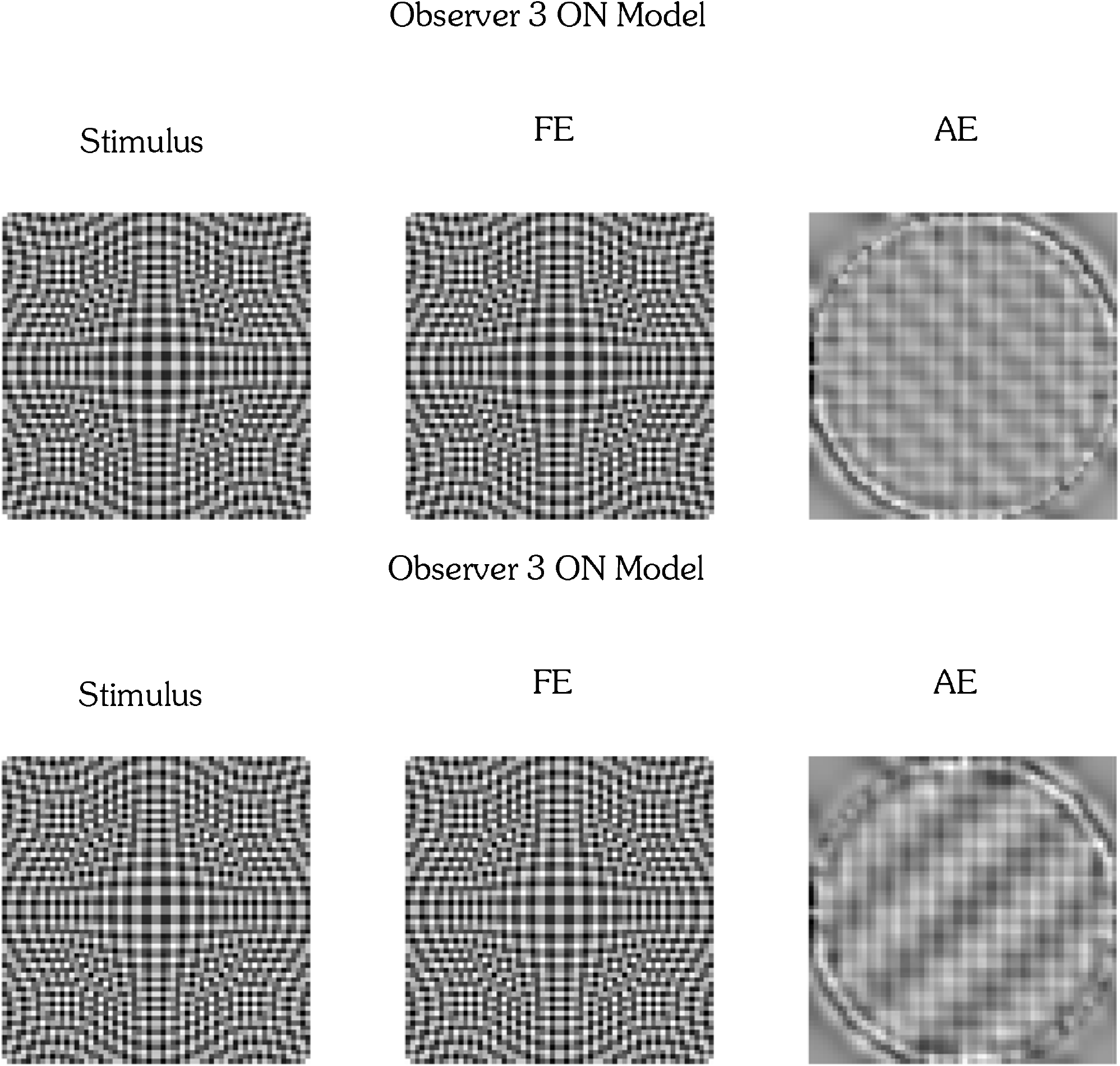
3D Shape-from-Texture stimuli with reconstructions generated by the FE and AE models. Predictions for Obs 3.

It took a combination of perceptual perceptogram measurements and generative computational modeling to reveal how neuronal RF properties lead to veridical and phantom percepts. Asymmetric matching has long been used as a psychophysical tool by putting the two eyes in different adaptation states, but the measured temporary differences are small and not as dramatic as the stable differences between the perceptograms and the viewed stimuli, which are due to signals from the two eyes going through different cortical filters. Invasive methods for measuring fine-grained neuronal properties such as electrophysiology and 2-photon imaging cannot be done in humans, and it remains to be seen if recording and perceptography^3^ in non-human primates can be expanded to a large enough scale. Non-invasive techniques such as fMRI and VEP would have to improve considerably in spatial and temporal resolution to answer such questions. We are exploring making more efficient and general Generative Phenomenology procedures by using generative adversarial networks^46^ to search through human similarity space for plaid or broader classes of patterns^47^ for the most probable perceptogram choice on each trial, with the observer’s responses serving as part of the adversarial network. This will be particularly helpful in developmental studies of form perception with children. As a result, for the foreseeable future, Generative Phenomenology could be a powerful strategy to understand the neural bases of human perception.

## MATERIALS AND METHODS

### Observers

Five female observers with amblyopia participated in the study. The gender distribution was due to chance and not due to selective recruitment. Each had a prior clinical diagnosis of amblyopia and received a thorough optometric evaluation by the authors J.W. and B.R. prior to the experiment. Of the group, two had strabismic amblyopia and three had anisometropic amblyopia. Detailed participant data are presented in Table 1. All participants were unaware of the study’s purpose and wore their best optical correction for the testing distance during the experiments. All participants provided written informed consent, and the study protocol was approved by the Institutional Review Board at the SUNY College of Optometry, in accordance with the tenets of the Declaration of Helsinki.

### Perceptograms

#### Apparatus

Figures 1C and 1D show the visual stimuli — test stimulus outlined in red and measuring stimuli outlined in blue — as presented using a Planar SD2620W Stereo/3D system. This setup consists of two calibrated LCD monitors positioned at right angles and optically combined using a beam splitter (Figure 1B). Each LCD monitor emits plane-polarized light, and the two monitor’s polarization axes were mutually orthogonal. The image in the beam splitter is viewed through glasses polarized so each eye receives input from only one monitor: the amblyopic eye (AE) views the test stimulus, and the fellow eye (FE) views the measuring stimuli. Prior to the experiment, we confirmed the absence of crosstalk between the two eyes and ensured that all nine patterns appeared evenly spaced. For one participant (Obs 4) with strong interocular suppression, the test and motion stimuli were alternated at a rate controlled by the observer. Each eye received a display resolution of 1920 × 1200 pixels at a 60 Hz refresh rate. Stimuli were presented and the experiment was controlled using PsychoPy, and all data were analyzed using Python. Head position was stabilized with a chin rest at a viewing distance of 1.2 meters. All experiments were conducted in a darkened room.

#### Stimulus generation

Using Python, we generated 24 test gratings that varied in orientation (0°, 45°, 90°, and 135°), spatial frequency (6, 9, and 12 cycles/deg), and stimulus polarity (ON: black background making bright bars salient; OFF: white background making dark bars salient). Each stimulus subtended 3° of visual angle. Phantom patterns were created by summing pairs of gratings, as illustrated in Figure 1.

#### Psychophysical procedures

At the start of the experiment, we confirmed that all the participants could perceive a veridical grating through the fellow eye (FE) by asking them to verbally describe the single grating. In the main task, the amblyopic eye (AE) viewed a central single grating (red outline), while the FE was presented with a surrounding test grating and seven plaid patterns, each representing a different phantom type (blue outlines), as shown in Figure 1C. The observer selected the surrounding pattern that most closely matched the central grating. As illustrated in Figure 1D, they then fine-tuned four parameters each of the component gratings of the selected plaid to improve the perceptual match. After adjustment, the observer judged whether the single grating viewed through the AE perceptually matched the plaid viewed through the FE. Gratings of the 4 orientations, 2 stimulus polarities, and 3 spatial frequencies, were presented in randomized order. Each session lasted approximately one hour, and the experiment was repeated on three separate days. If the perceptogram did not provide a perfect match, the observer first provided a verbal description of the mismatch. When verbal reports were insufficient, the observer sketched the perceived phantom using Procreate on a 12.9-inch iPad. The drawing was projected onto the Planar display seen by the FE, while the AE continued viewing the single grating.

### Reverse Correlation

#### Stimulus generation

To measure orientation tuning, we presented a rapid sequence of sinusoidal gratings randomly varying in orientation at 10 Hz, with each grating displayed for 100 ms. This temporal frequency was chosen for task feasibility, as higher frequencies were too difficult for some participants with amblyopia. Gratings had a spatial frequency of 6.0 cycles/deg— the lowest frequency used in the Perceptogram experiment—and were shown at full contrast. Stimuli were presented on the front-facing monitor of the Planar Display, without the beam splitter, centrally through a 3° diameter circular aperture, against either a white or black background. Each condition consisted of one of four target orientations (0°, 45°, 90°, or 135°) combined with one of the two stimulus polarities (ON or OFF) and was tested separately for the fellow and amblyopic eyes, yielding a total of 16 unique conditions. Each trial within a condition consisted of 600 gratings presented over 60 seconds. Gratings were randomly selected from 10 possible orientations (spaced 18° apart) and one of four phases (0, π/2, π, or 3π/2).

#### Psychophysical procedures

The experimental procedure followed protocols established in previous studies^17-20^. To assess each eye independently, participants wore an eye patch over the non-tested eye. Each trial consisted of a 1-minute sequence during which participants were instructed to press a button as quickly as possible upon detecting a grating with a specific orientation (horizontal, vertical, or oblique) embedded within the sequence. Trials began with brief on-screen instructions, followed by a brief tone signaling the onset of the grating sequence. Participants initiated each trial with a button press and were instructed to maintain central fixation throughout. After each trial, the instruction screen reappeared, allowing participants to rest before continuing. No performance feedback was provided. Each condition included 30 trials, lasting approximately 45 minutes in total. The full set of conditions—four orientations, two stimulus polarities (ON and OFF), and two eyes (fellow and amblyopic)— was administered in randomized order, typically with two conditions completed per day.

### Mathematical & Statistical Details

#### Cortical model

In this section, we provide the equations that describe how we built filters for FE and AE, and how we tested the adequacy of the filters to reconstruct the perceptograms. The FE filters are standard, but we describe details to be able to specify what was transformed for AE. Because the fellow eye (FE) did not exhibit any distortions, we assume that signals are processed like normal V1, and we model it with a steerable pyramid for frequency decomposition. The Steerable Pyramid python library is publicly available at https://docs.plenoptic.org/docs/branch/main/tutorials/models/Steerable_Pyramid.html. The steerable pyramid decomposition operates in the Fourier domain. Let *I* (*x,y*) be the input image. Its 2D Fourier transform is ℱ (*I* (*x,y*)), and the inverse Fourier transform ℱ^−1^ (ℱ *(I* (*x,y*))) reconstructs the original image from its frequency representation. Figure 2A illustrates the architecture of the complex steerable pyramid. The left side of the block diagram depicts the decomposition process, while the right side illustrates reconstruction. The decomposition is performed in polar coordinates, where the radial component *r* represents spatial frequency and the angular component *θ* corresponds to orientation. The high-pass and low-pass filters used in this transform are defined as follows:

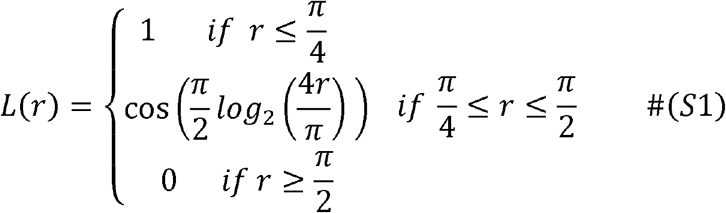

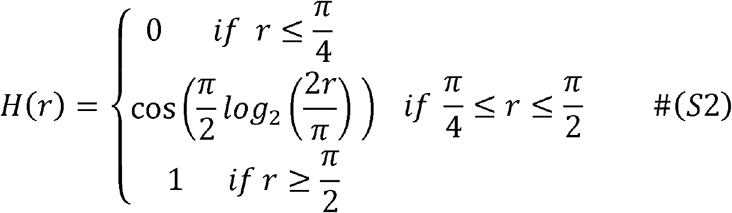

The input image is first decomposed into high-pass and low-pass components by multiplying its Fourier transform with the filters, *H*_0_ and *L*_0_

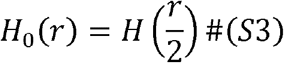

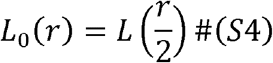

The low-pass band is subsequently decomposed into a lower-frequency band and a set of orientation-selective sub-bands through the combination of a high-pass filter and an angular mask (*G*_*k*_(*θ*)):

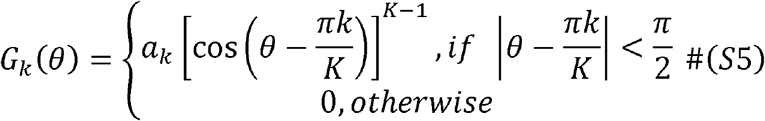

Here, *K* the preferred is the total number of orientation-selective filters, *K* indicates the preferred orientation, and *a*_*k*_ is a normalization constant.

The orientation symmetric filer in the first scale, *B*_0,*k*_ is expressed as:

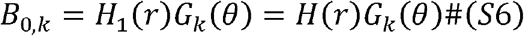

Following this cascade, *L*_0_ is first applied to isolate the low-frequency band, and then multiplied by *B*_0,*k*_ to create an oriented band-pass filter that is selective for both spatial frequency and orientation. To proceed to the next scale (i.e., lower spatial frequency), the model applies *L*_1_ (= *L*) and repeats the process by multiplying it with the orientation-symmetric filter *B*_1,*k*_ (=*H*_2_*G*_*k*_=*H*(2*r*)*G*_*k*_). The filter response is zero beyond ∣r∣>π/2, allowing computations to be limited to ∣r∣<π/2 for efficiency. When filters across scales are combined, they are zero-padded to ensure matching dimensions before summation. This process is repeated across the specified number of scales.

In general, the orientation-symmetric filter at the *i*-th scale and *k*-th orientation, denoted *B*_*i,k*_, is defined as:

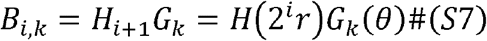

*i*-th scale lowpass filter *L*_*i*_(*r*) is *L*(2^*i*−1^*r*). Therefore, the oriented band-pass filter at *n*-th scale and *k*-th orientation is constructed as:

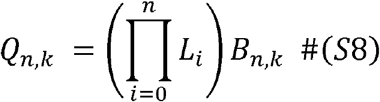

To visualize the spatial profile of these filters, a centered delta function *δ* (*x,y*) is passed through each filter. The resulting spatial domain representation is obtained by:

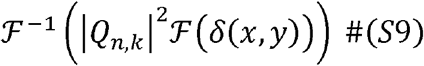

These visualizations are shown in Figure 2A, where four spatial scales are arranged from top to bottom (from high to low frequency), each with 16 orientations. The responses of these filters satisfy perfect reconstruction under the *L*^2^ -norm.

Figure S4B illustrates the architecture of the complex steerable pyramid for AE incorporating the Amblyopic linear transformation *A*_*i,k*_ for each *i*-th scale and *k*-th orientation filter. The model follows the same steps as in the FE V1 pathway, except that the output of each *B*_*i,k*_ is transformed by *A*_*i,k*_. We estimated *A*_*i,k*_ as follows: Let ***S*** (∈ *R*^*HW×C*^) be a matrix containing vectorized single grating stimuli, where *H* and *W* are the image height and width, and *C* is the number of conditions. Similarly, let D (∈ *R*^*HW×C*^) represent the corresponding perceptograms. We assume the oriented band-pass filter response *Q*_*i,k*_ is subjected to a linear transform *A*_*i,k*_ (∈ *R*^*HW×HW*^), yielding the AE cortical response:

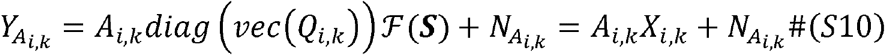

Where 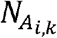, represents an independent noise source and *X*_*i,k=*_ *diag* (*vec* (*Q*_*i,k*_)) ℱ(***S***) The corresponding FE response to the perceptograms is:

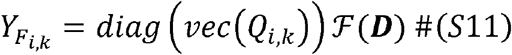

Assuming the linking hypothesis that 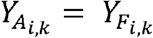, the cross-spectrum between 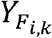 and *X*_*i,k*_ becomes:

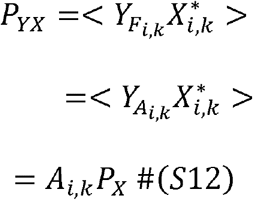

Where 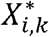 is the conjugate transpose of *X*_*i,k*_ and 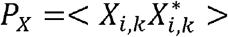 Then, the amblyopic transformation *A*_*i,k*_ can be estimated as:

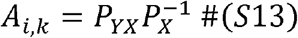

The AE V1 filters for Obs 3 for the ON condition are visualized in Figure 2B using the same method as Equation S9.

For each observer, session, and stimulus type (ON vs. OFF), we tested whether the discrete set of filters for that condition could adequately reconstruct the perceptograms. For *S* (*x, y*) a grating from the stimulus set, and *D* (*x,y*) the corresponding perceptogram, we passed the Fourier transform of the grating through the appropriate set of AE filters, and then compared the inverse Fourier transform which gave the reconstruction to the perceptogram:

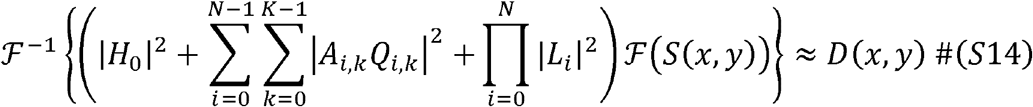

### Sinusoidal Deformations from Circularity

This section describes the autocorrelations computed for the simulations. Individuals with amblyopia exhibit a deficit in detecting sinusoidal modulations from circularity of the fourth derivative of a Gaussian (D4)^15^. Using similar procedures as above, we computed FE and AE filter responses for the D4 reference and various amplitudes for each frequency of modulation. The image was reconstructed using Equation S14, followed by computation of its rotational correlation. Each rotational autocorrelation was computed as:

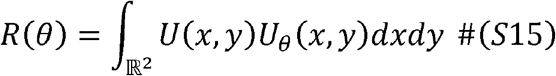

*U* (*x, y*) is the reconstructed image by the methods above and *U*_*θ*_ is (*x, y*) rotated by angle *θ* around the origin. We sampled rotation angles from −17° to 17° in 1° increments, yielding a vector of length 35. Cosine similarity between the auto-correlation vectors of the reference and modulated shapes was computed to generate the plots shown in Figure 4C.

### Orientation Tuning via Reverse Correlation

This section provides details of the data analysis of the reverse correlation experiment. In the Reverse Correlation Experiment, observers fixated on the central display and were instructed to press a button as soon as they detected the target orientation. Grating orientations presented during the 1-second window (10 frames) prior to each key press were collected and used to construct a histogram of occurrences across time (τ) and orientation (*θ*). The concatenated histograms were combined into a heatmap as a function of orientation (*θ*) and time (τ). Each trial consisted of a 600-frame (60-second) time series, which was resampled 5,000 times using bootstrap resampling. To reduce noise, the heatmap was smoothed using a small symmetric Gaussian kernel with *σ*_*θ*_ = 6 ° and *σ* _*τ*_ = 0.033 seconds. The resulting heatmap was then normalized to produce PF(*θ, τ*). Individual PF(*θ, τ*) were computed for the Non-Distortion and Distortion Groups. As the individual trends within each group were consistent, we averaged the data across two observers for the Non-Distortion Group and three observers for the Distortion Group. To examine orientation tuning more closely, we marginalized the PF(*θ, τ*) over *τ* to obtain a tuning curve. The resulting data were then fit with an additive von Mises function:

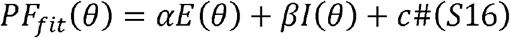

*E*(*θ*) and *E* (*θ*) denote the excitatory and inhibitory components of the orientation tuning curve, respectively. The parameters *α, β*, and *c* modulate the amplitudes of excitation and inhibition, as well as the baseline level of the response. Both *E*(*θ*) and *I*(*θ*) are defined using normalized von Mises functions:

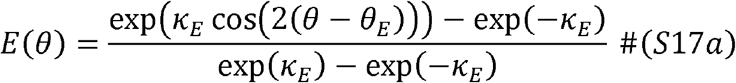

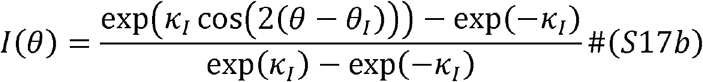

Here, *κ*_*E*_ and *κ*_*I*_ determine the tuning widths of the excitatory and inhibitory components, while *θ*_*E*_ and *θ*_*I*_ represent their respective preferred orientations.

## Supporting information

https://drive.google.com/file/d/1Rl25qFsH4PAUHuc0O6cNY-GCe7dEYvx7/view?usp=drive_link

## Acknowledgments

We thank Anya Hurlbert for suggesting the title.

## Author Contributions

Conceptualization: QZ, AM

Methodology: AM, QZ

Observer recruitment and screening: JW, BR, AM

Code: AM

Analyses and models: AM, QZ

Investigation: AM, BR, FO, JW, JMA, QZ

Visualization: AM, QZ

Writing: AM, QZ

Review & editing: AM, BR, FO, JW, JMA, QZ

## Data and Materials Availability

All data needed to evaluate the conclusions in the paper are present in the paper and the Supplementary Materials. The complete set of perceptograms and their reconstructions are included in the Supplemental file. The computational code is posted at: https://drive.google.com/drive/folders/1noWwEE7W2AubAtolAY7tRQIYy71l83Ia

