## Supplementary material for "Phantom Forms in Amblyopic Vision and what they reveal about the Generative Brain": https://drive.google.com/file/d/1Rl25qFsH4PAUHuc0O6cNY-GCe7dEYvx7/view?usp=drive_link

**SUPPLEMENTAL MATERIALS:**

Complete set of Perceptograms for 3 observers and 3 sessions, along with reconstructions from the Amblyopic cortical models derived for each observer and session:

**Figure S1. Visual comparisons of the complete set of measured and generated Perceptograms:** In each panel, the left two columns display ON stimuli, and the right two OFF stimuli. Each row corresponds to a different stimulus orientation. Within each pair, the left image represents the observer's perceptual match, while the right image shows the corresponding reconstruction generated by the transformed AE cortex filters. In 200 out of 216 conditions, the generated plaids matched the perceived distortions. In the other 16 conditions, observers provided verbal descriptions of the mismatch. When verbal descriptions were insufficient, observers traced their percepts using Procreate on an iPad.

#### 6 cycle/deg (Obs 3 Session 1)

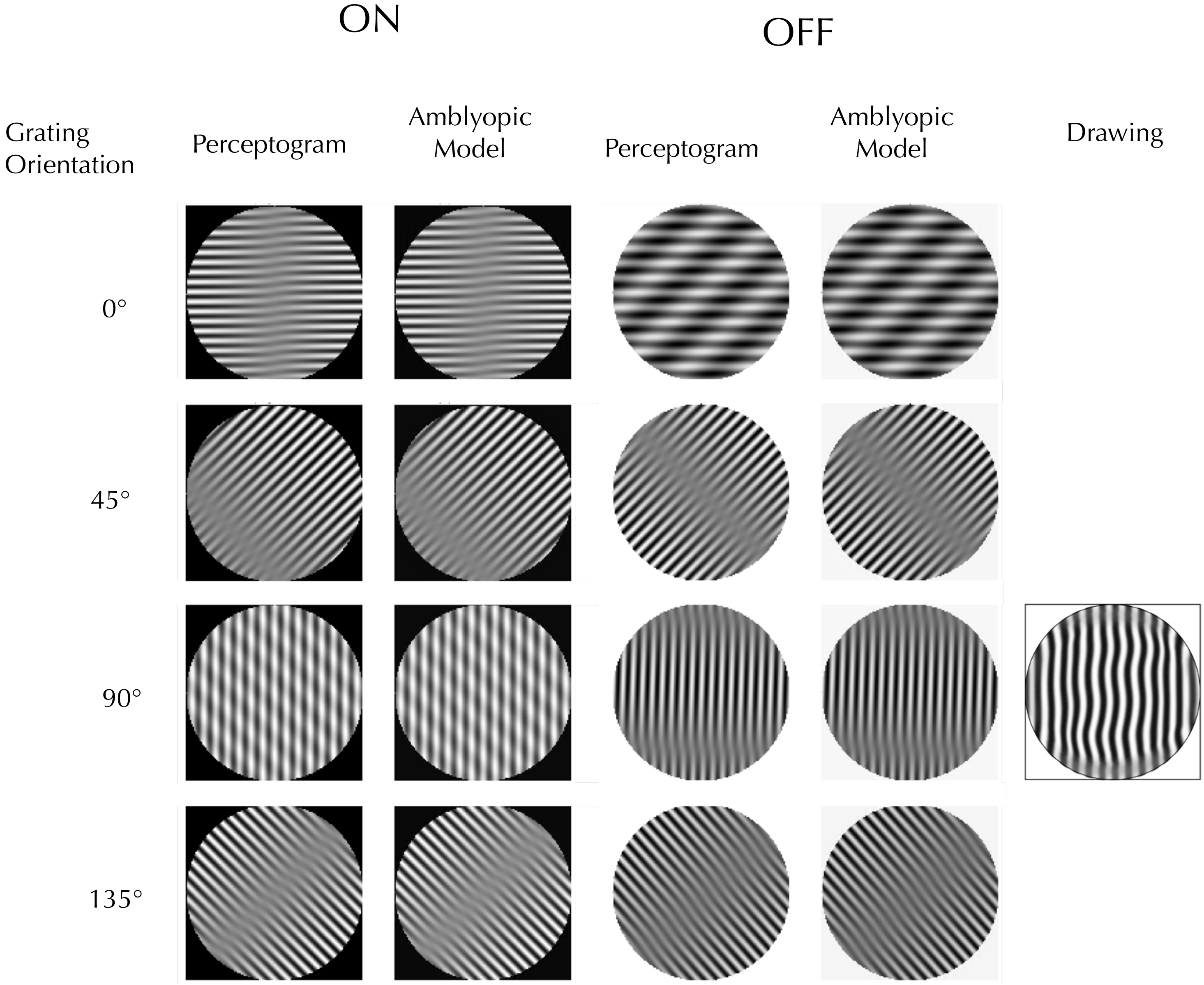

9 cycle/deg (Obs 3 Session 1)

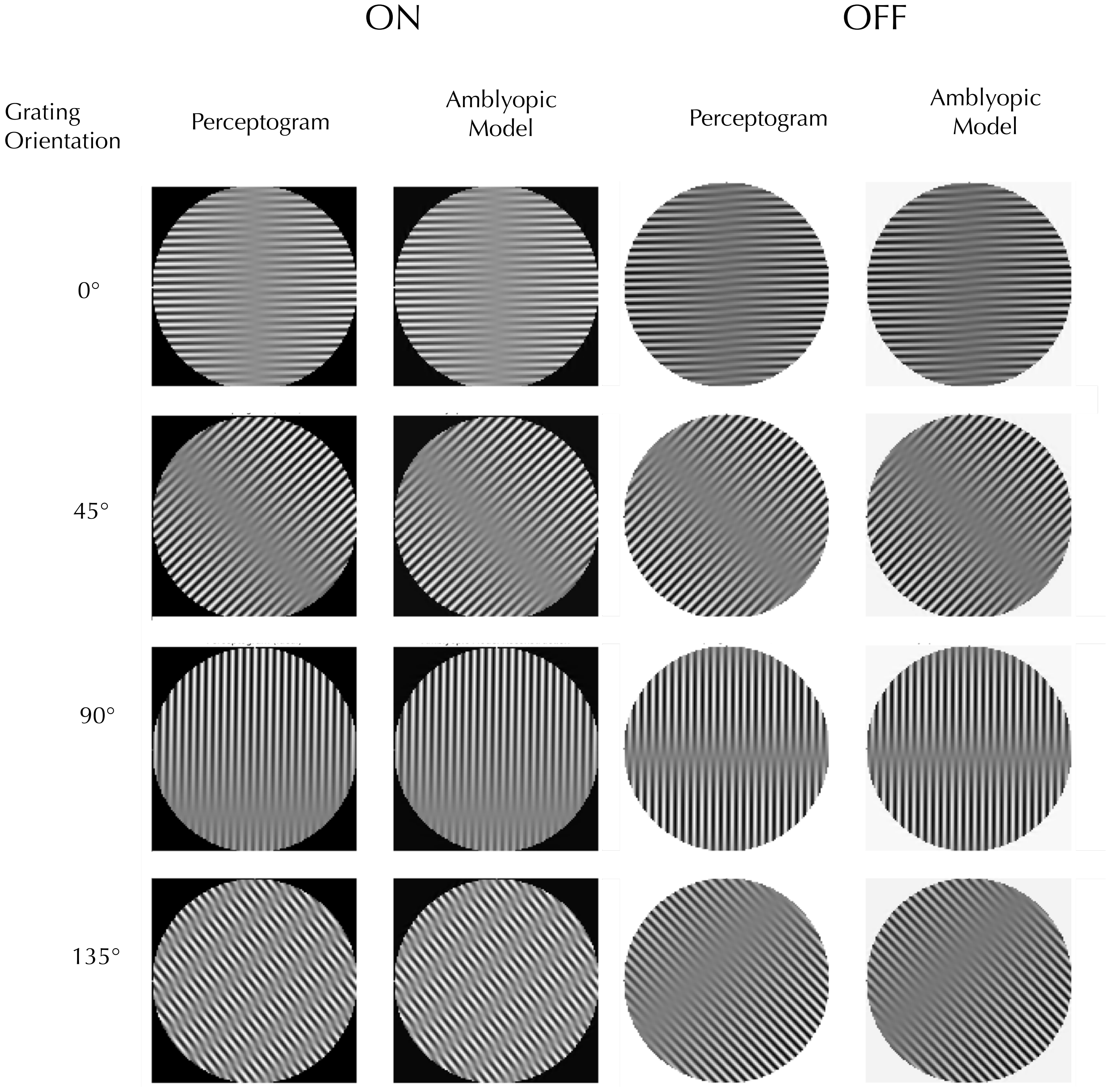

12 cycle/deg (Obs 3 Session 1)

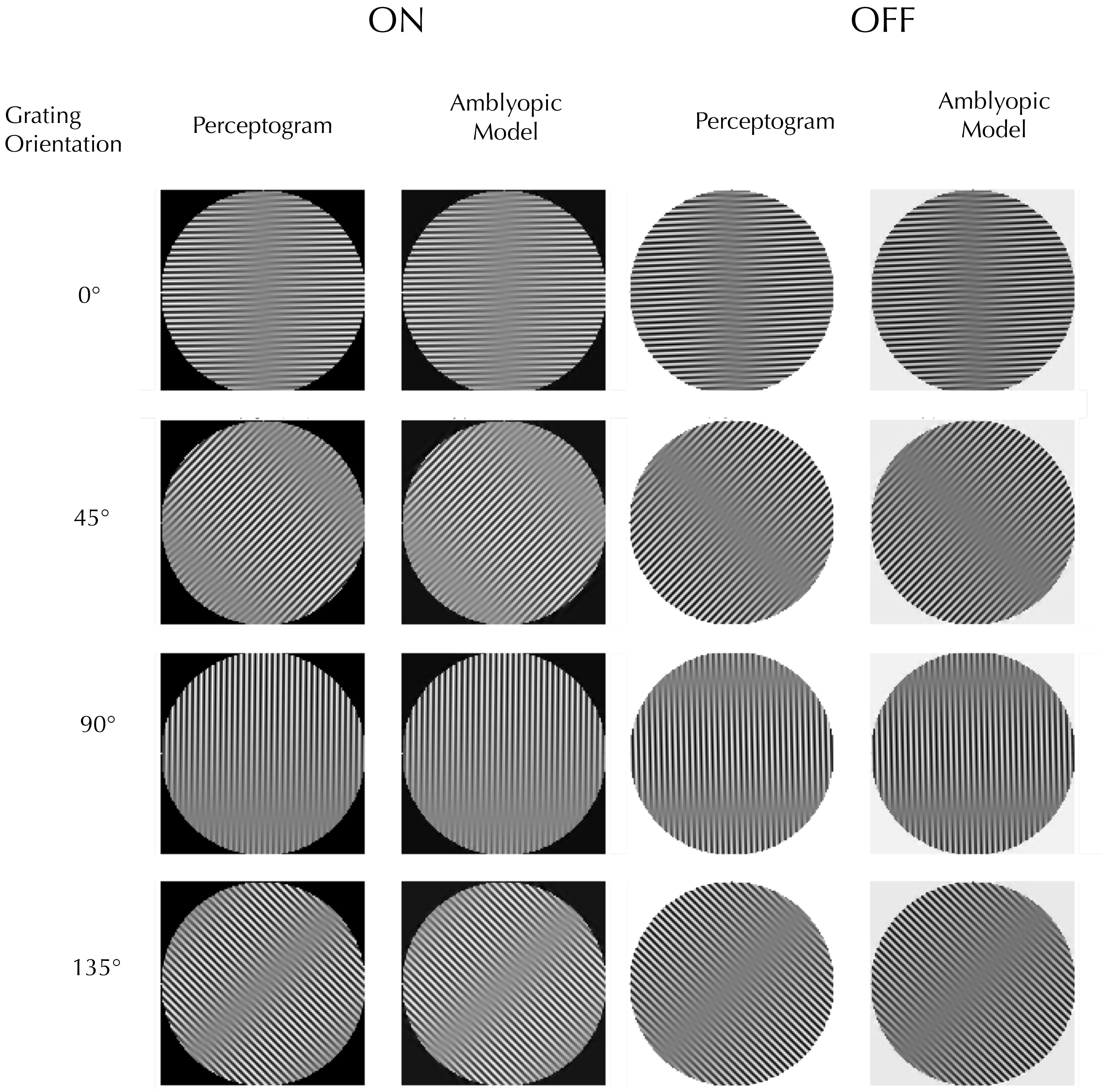

6 cycle/deg (Obs 3 Session 2)

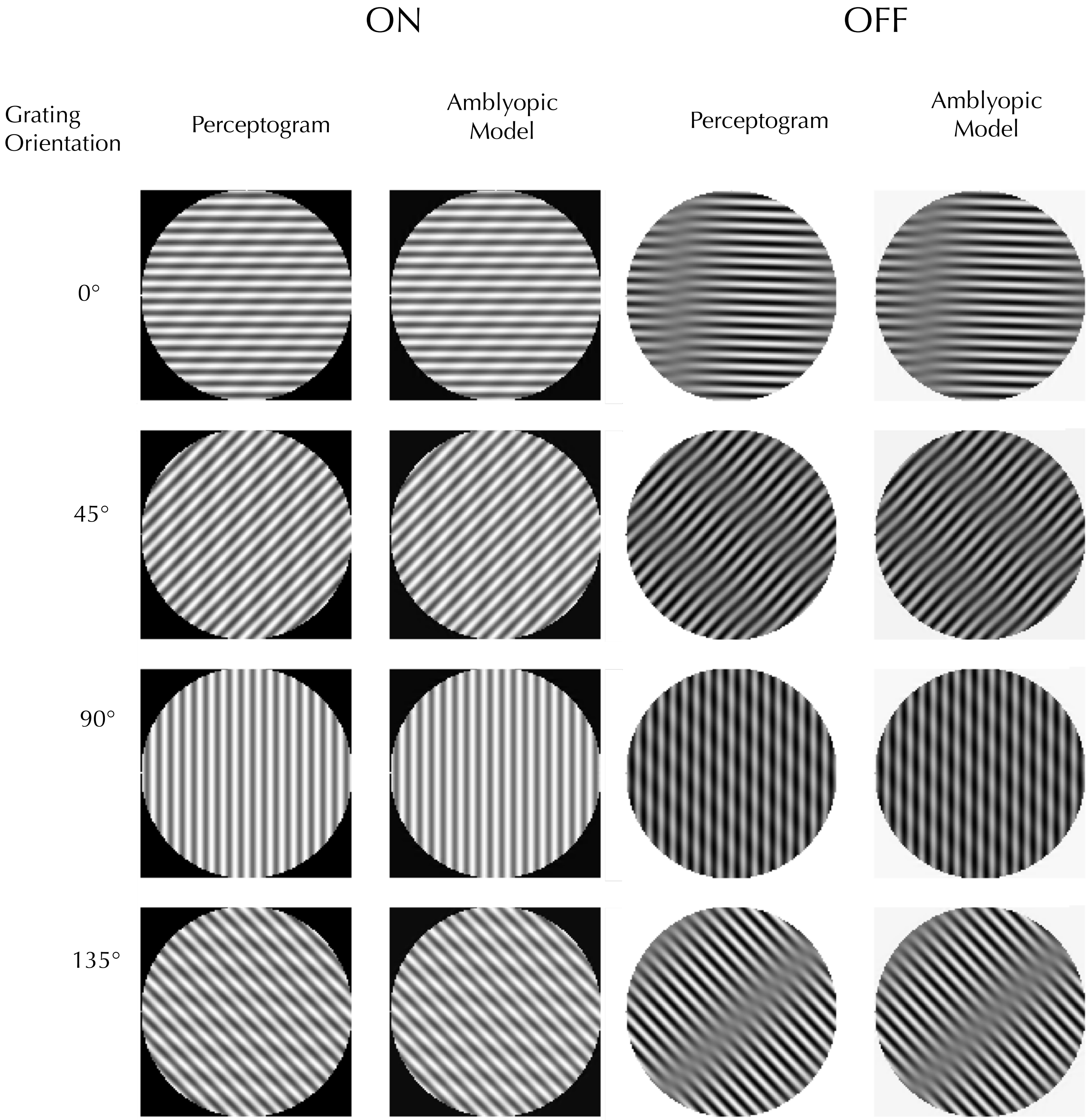

### 9 cycle/deg (Obs 3 Session 2)

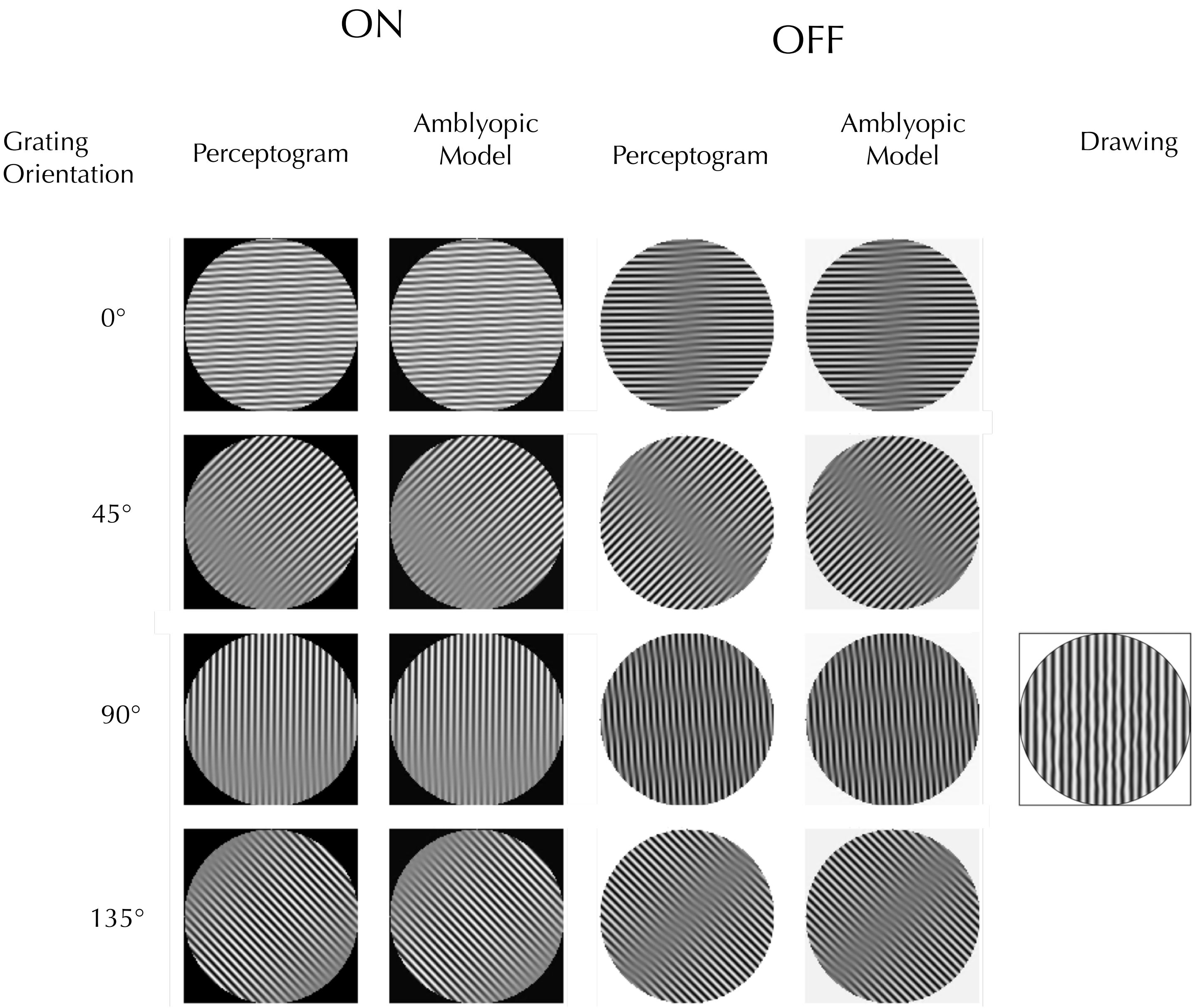

12 cycle/deg (Obs 3 Session 2)

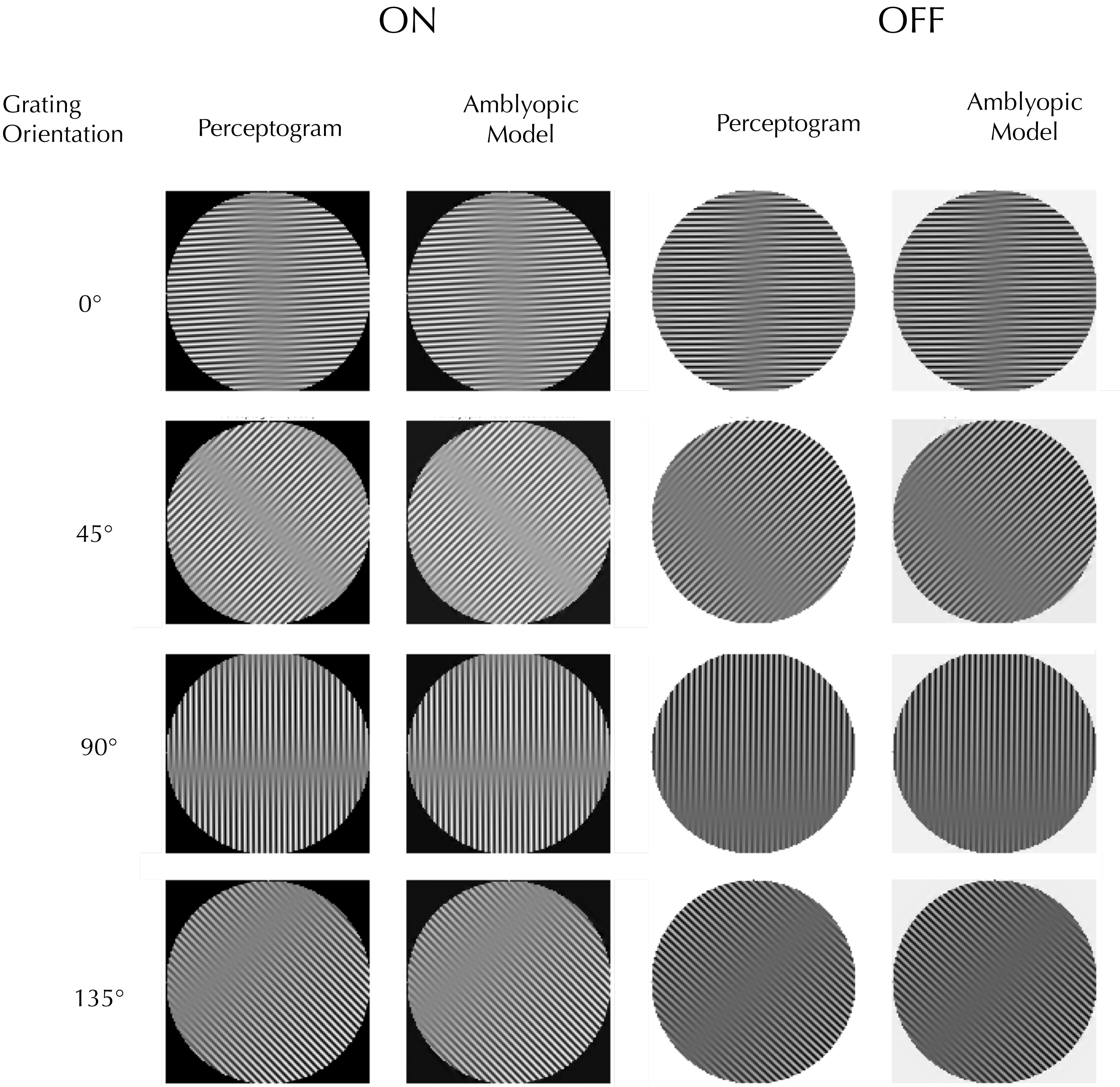

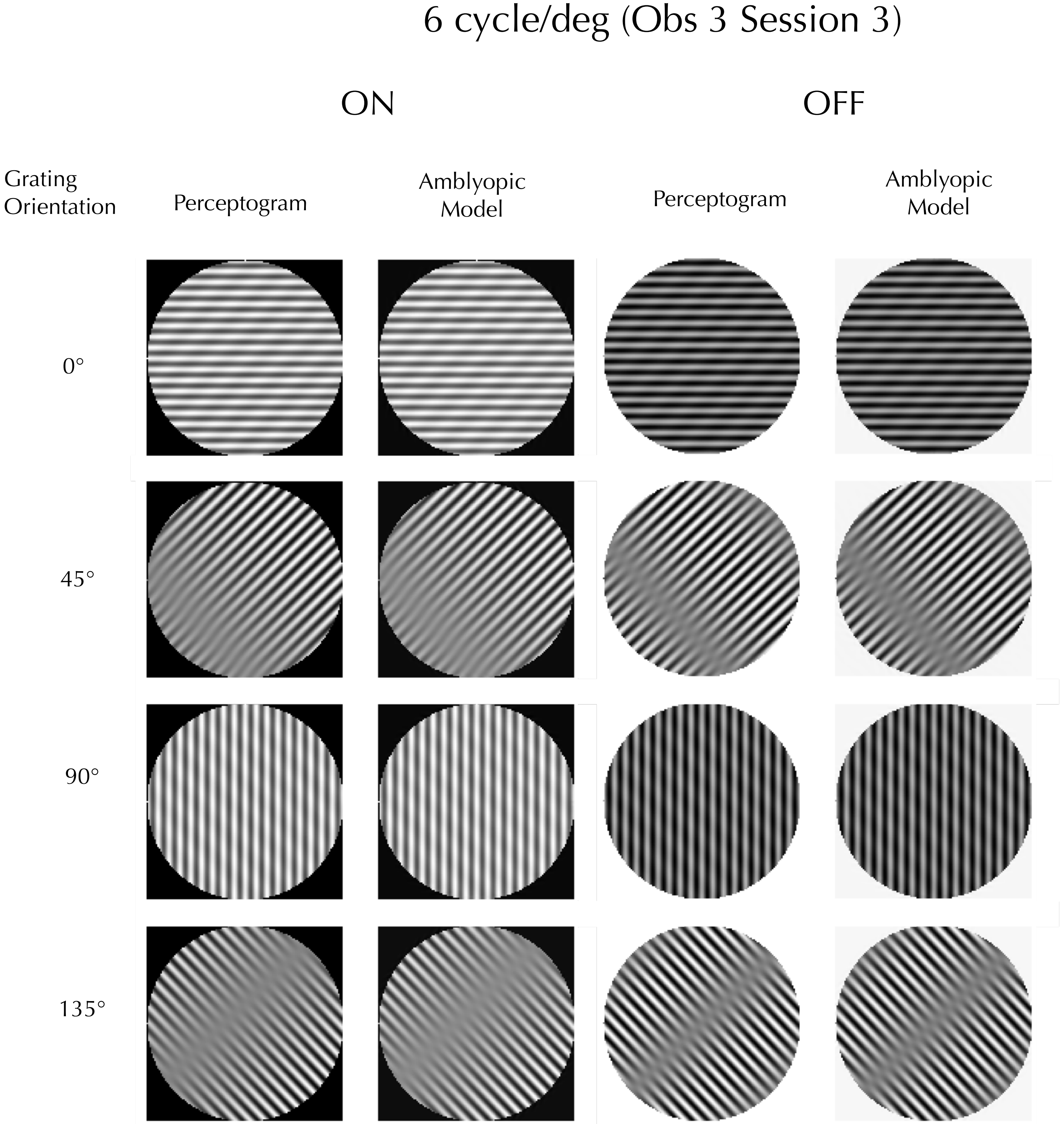
6 cycle/deg (Obs 3 Session 3)
ON
OFF
Grating Orientation
Perceptogram
Amblyopic Model
Perceptogram
Amblyopic Model
0°
45°
90°
135°

9 cycle/deg (Obs 3 Session 3)

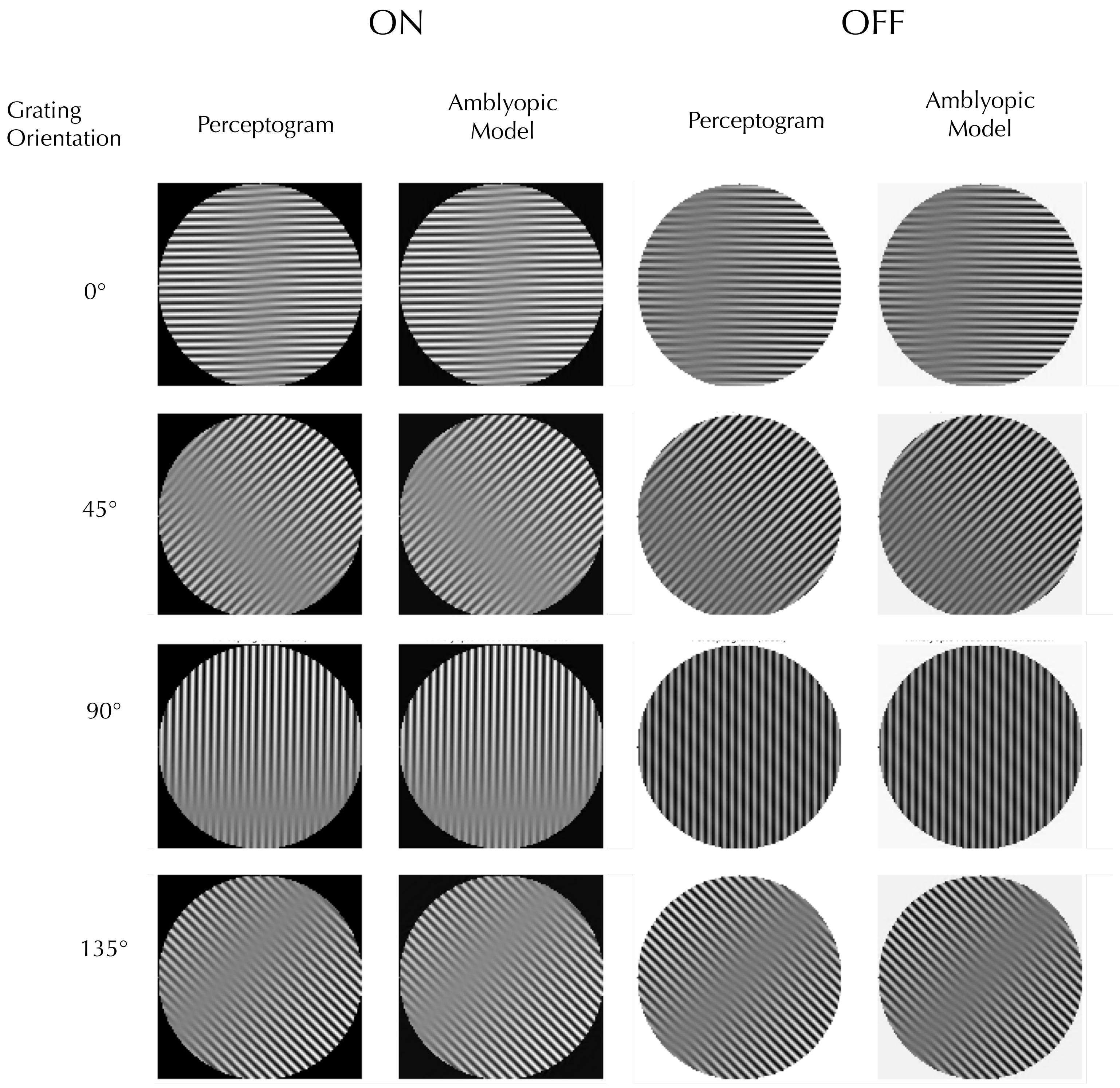

12 cycle/deg (Obs 3 Session 3)

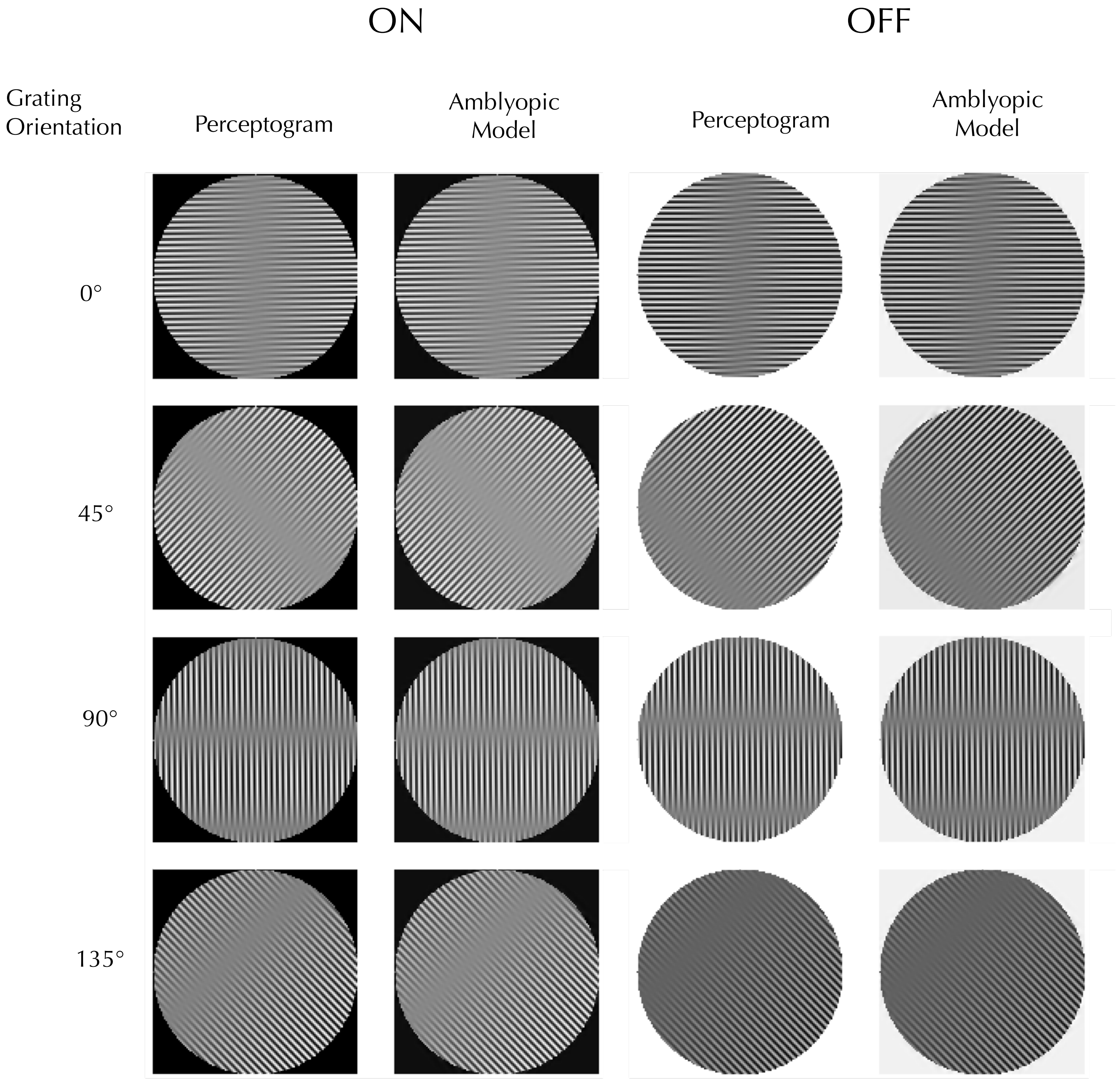

6 cycle/deg (Obs 4 Session 1)

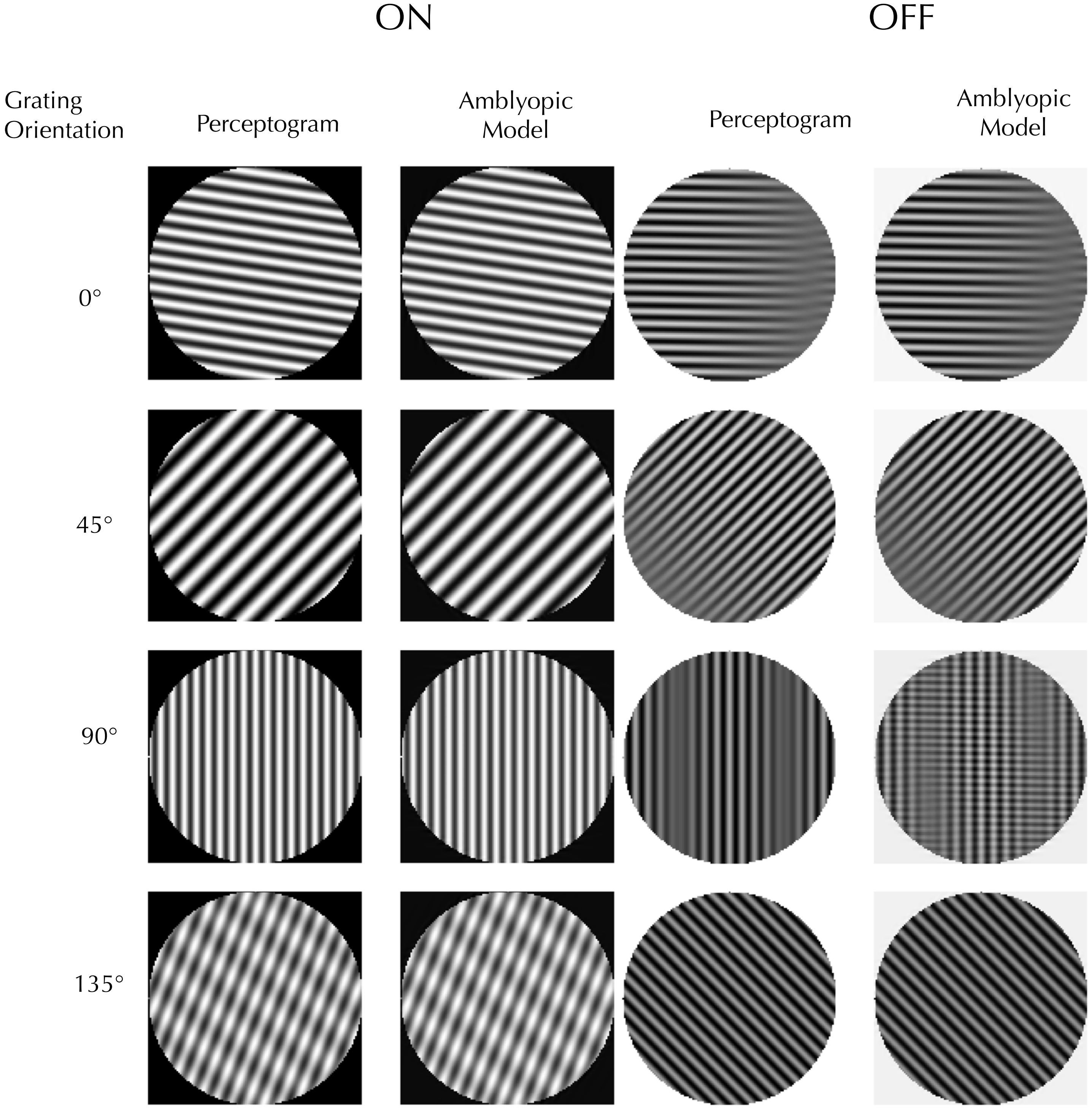

### 9 cycle/deg (Obs 4 Session 1)

ON

OFF

Grating Orientation | Perceptogram | Amblyopic Model | Perceptogram | Amblyopic Model | Verbal Description of Misalignment

0°

45°

90°

135°

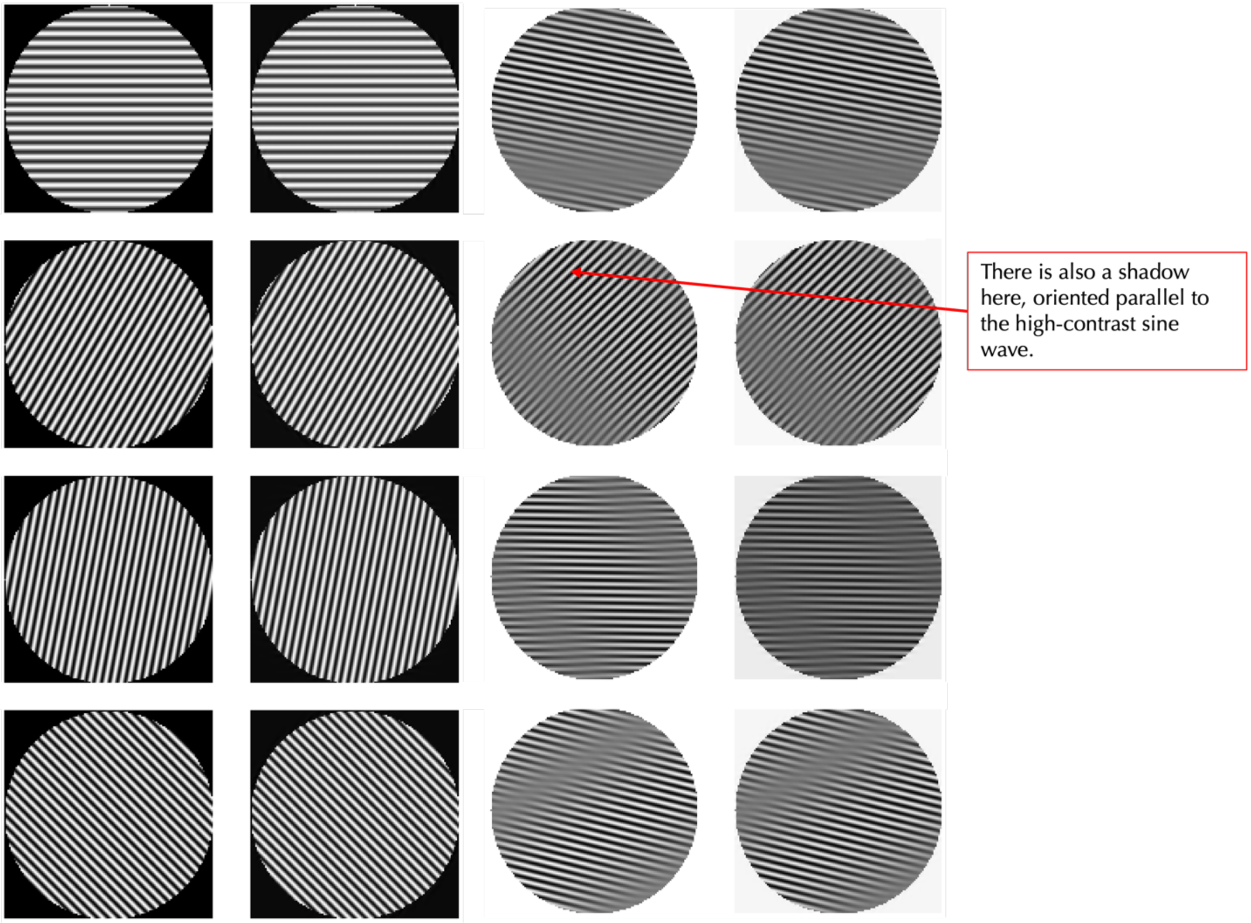

#### 6 cycle/deg (Obs 4 Session 2)

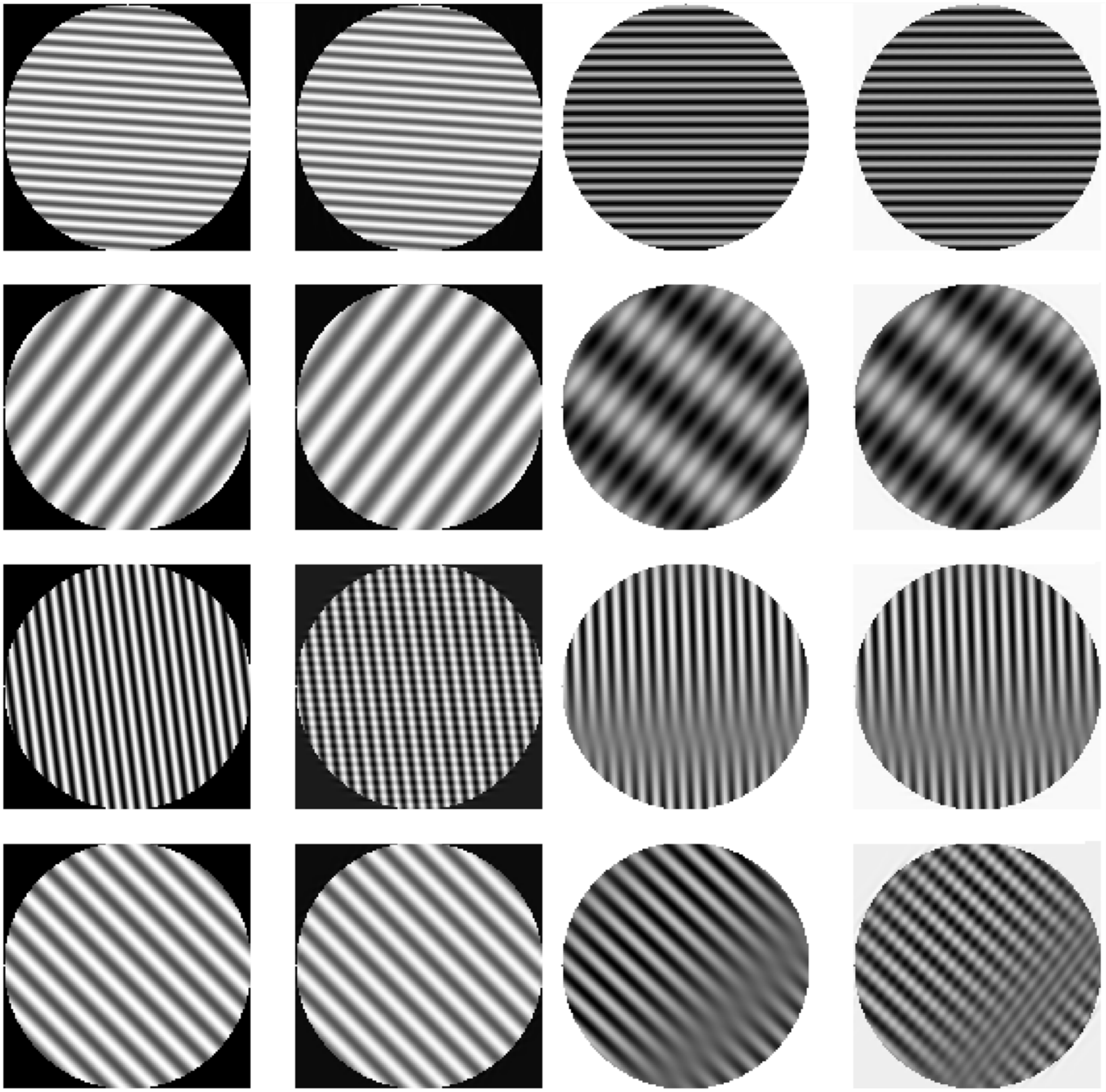

9 cycle/deg (Obs 4 Session 2)

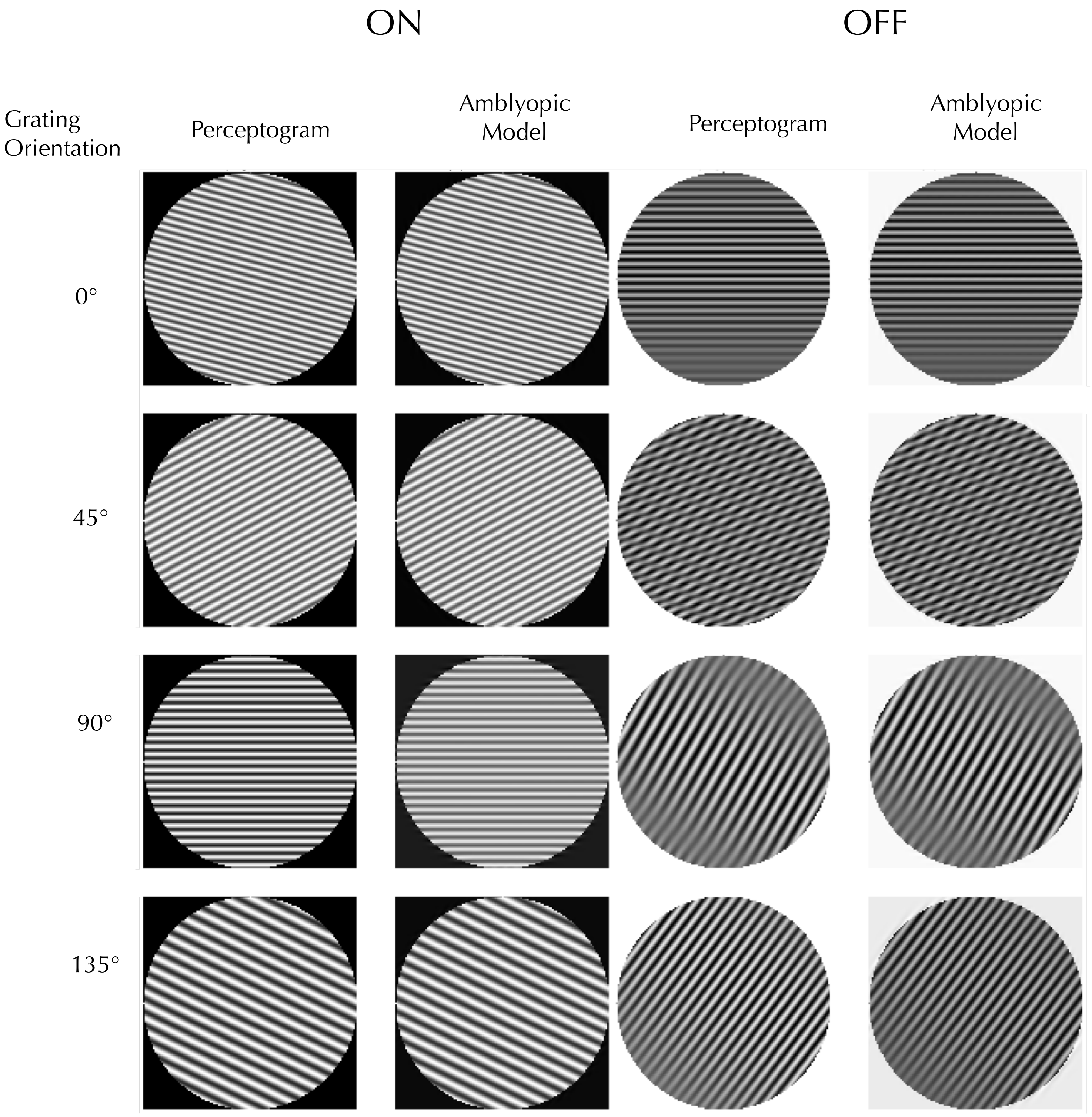

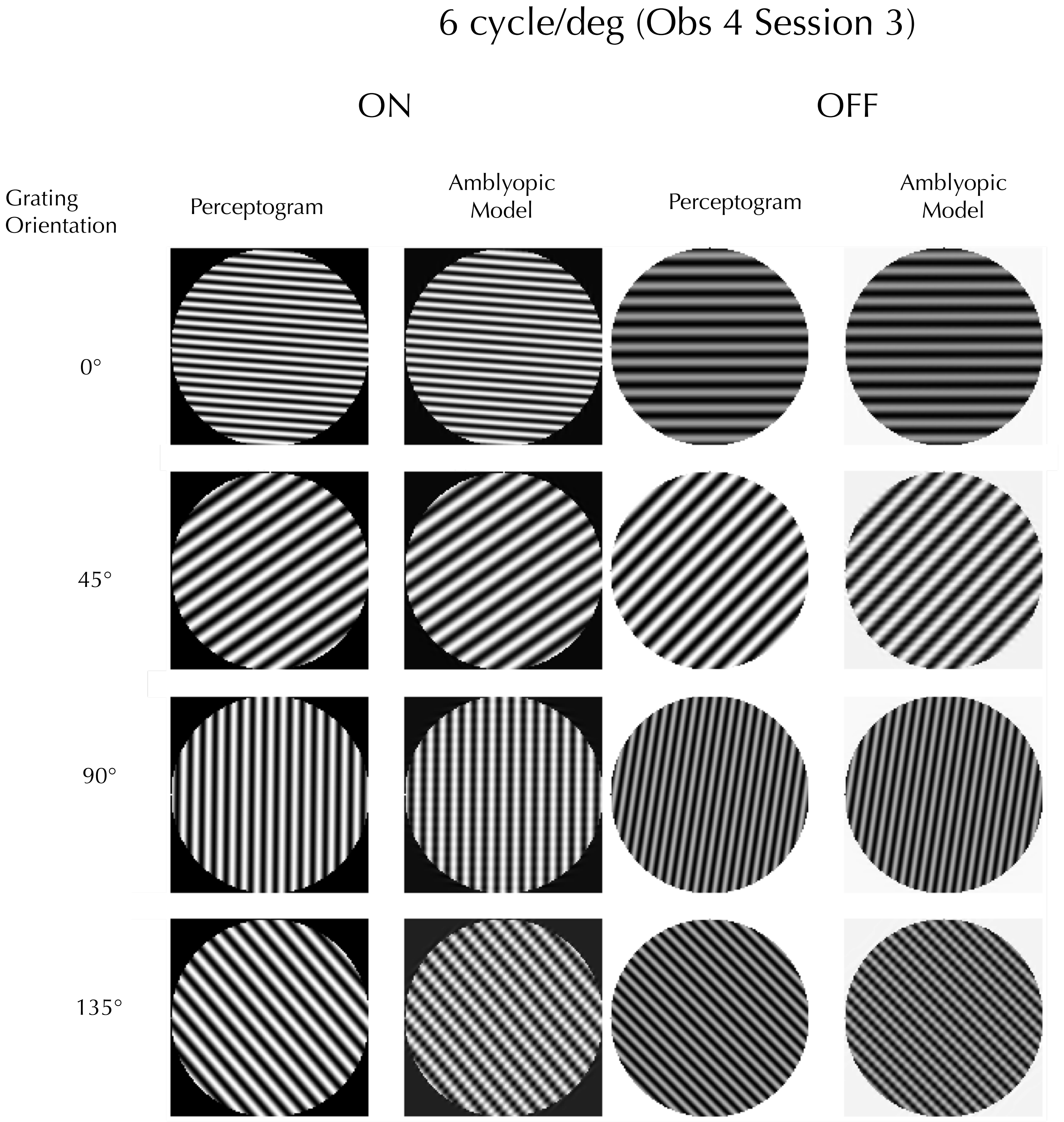
6 cycle/deg (Obs 4 Session 3)
ON
OFF
Grating Orientation
Perceptogram
Amblyopic Model
Perceptogram
Amblyopic Model
0°
45°
90°
135°

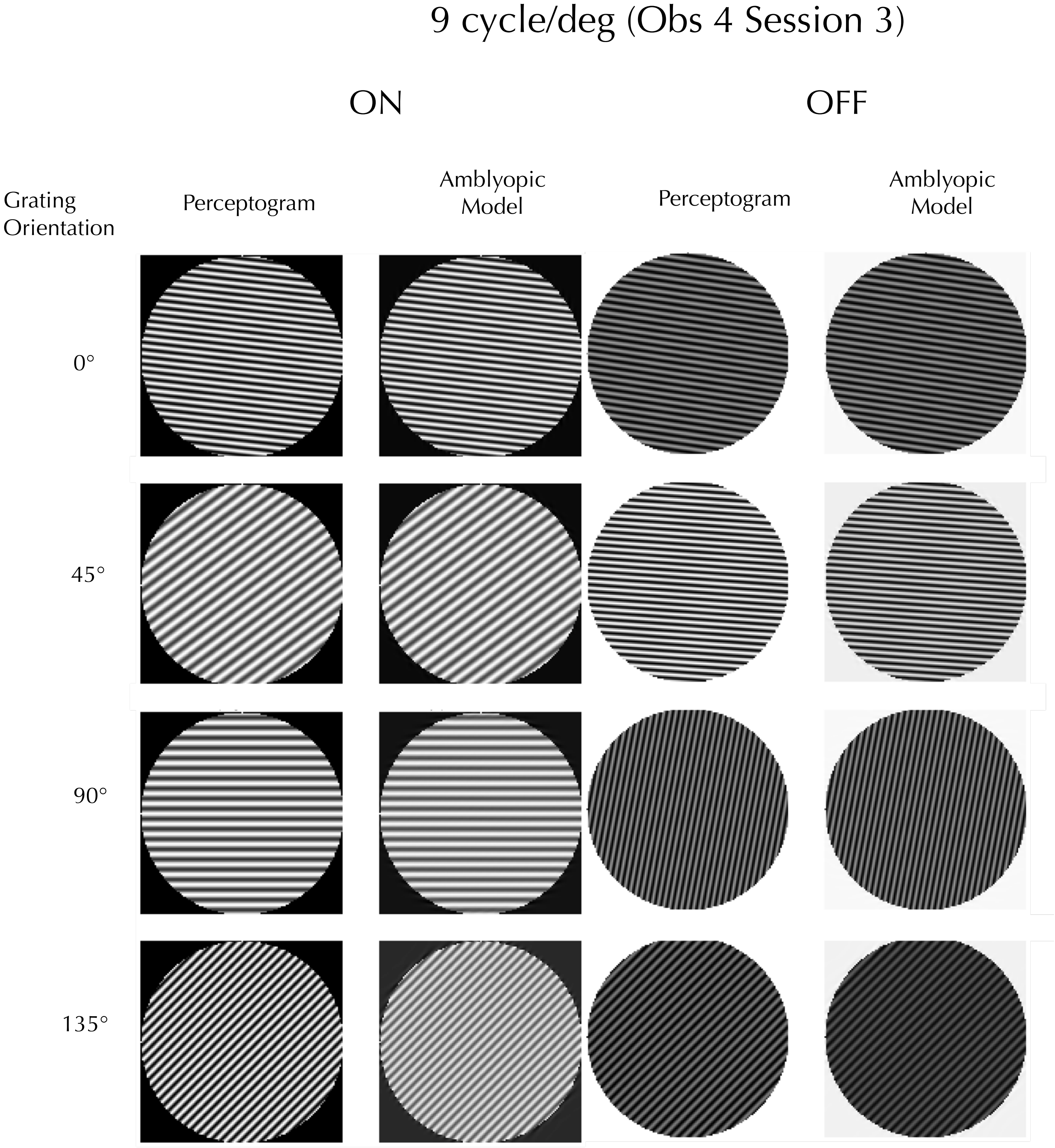
9 cycle/deg (Obs 4 Session 3)
ON
OFF
Grating Orientation
Perceptogram
Amblyopic Model
Perceptogram
Amblyopic Model
0°
45°
90°
135°

6 cycle/deg (Obs 5 Session 1)

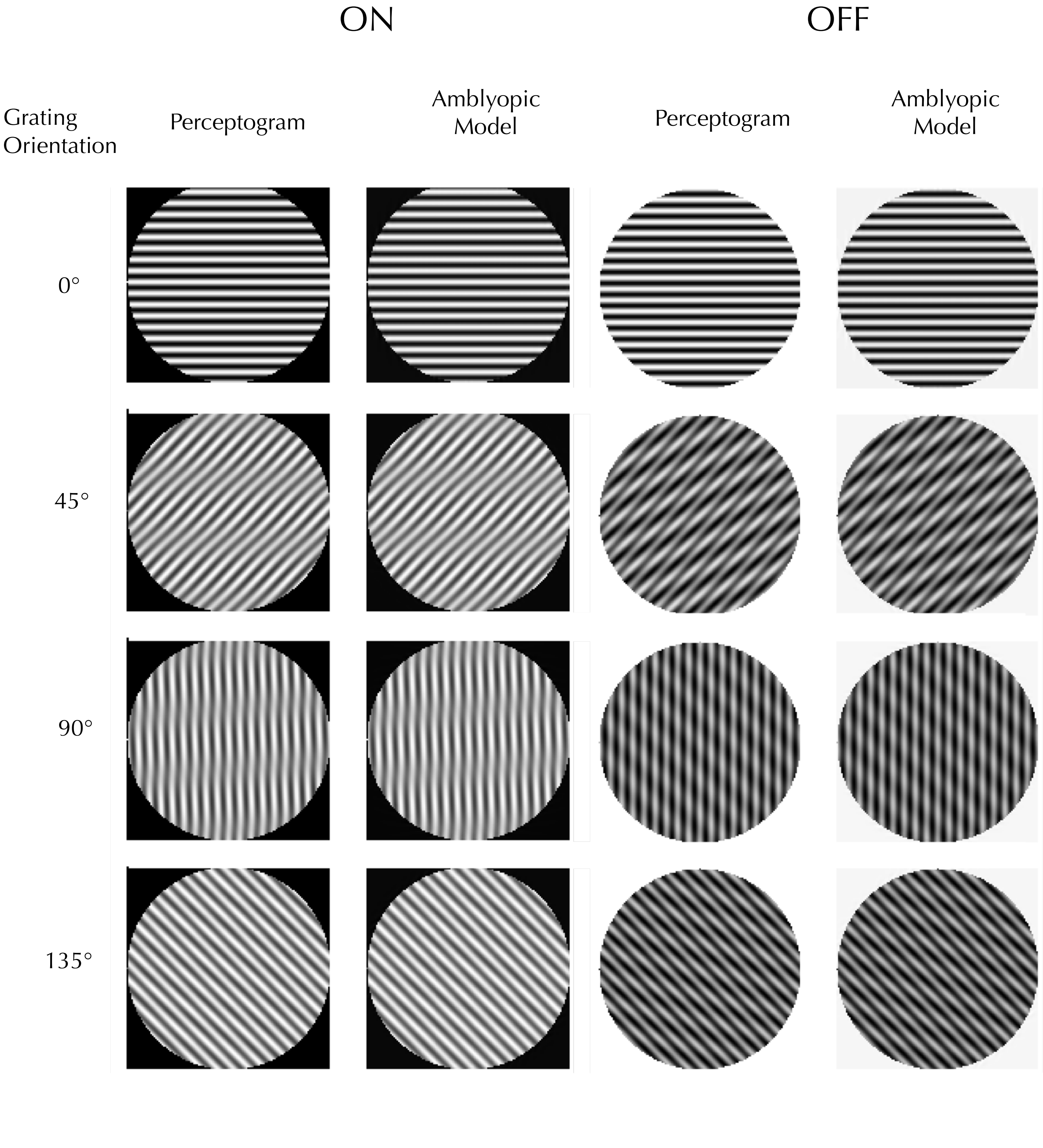

### 9 cycle/deg (Obs 5 Session 1)

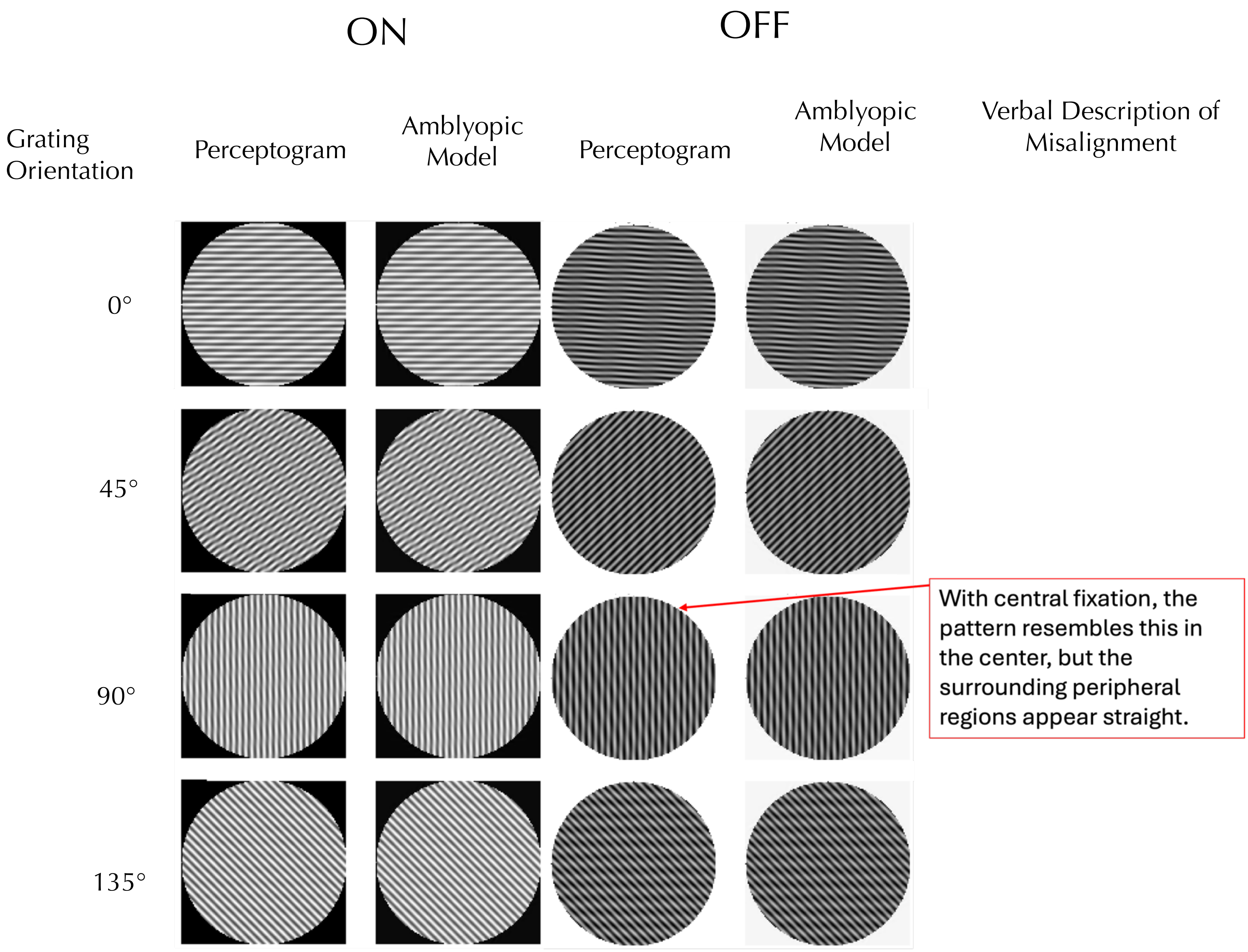

### 12 cycle/deg (Obs 5 Session 1)

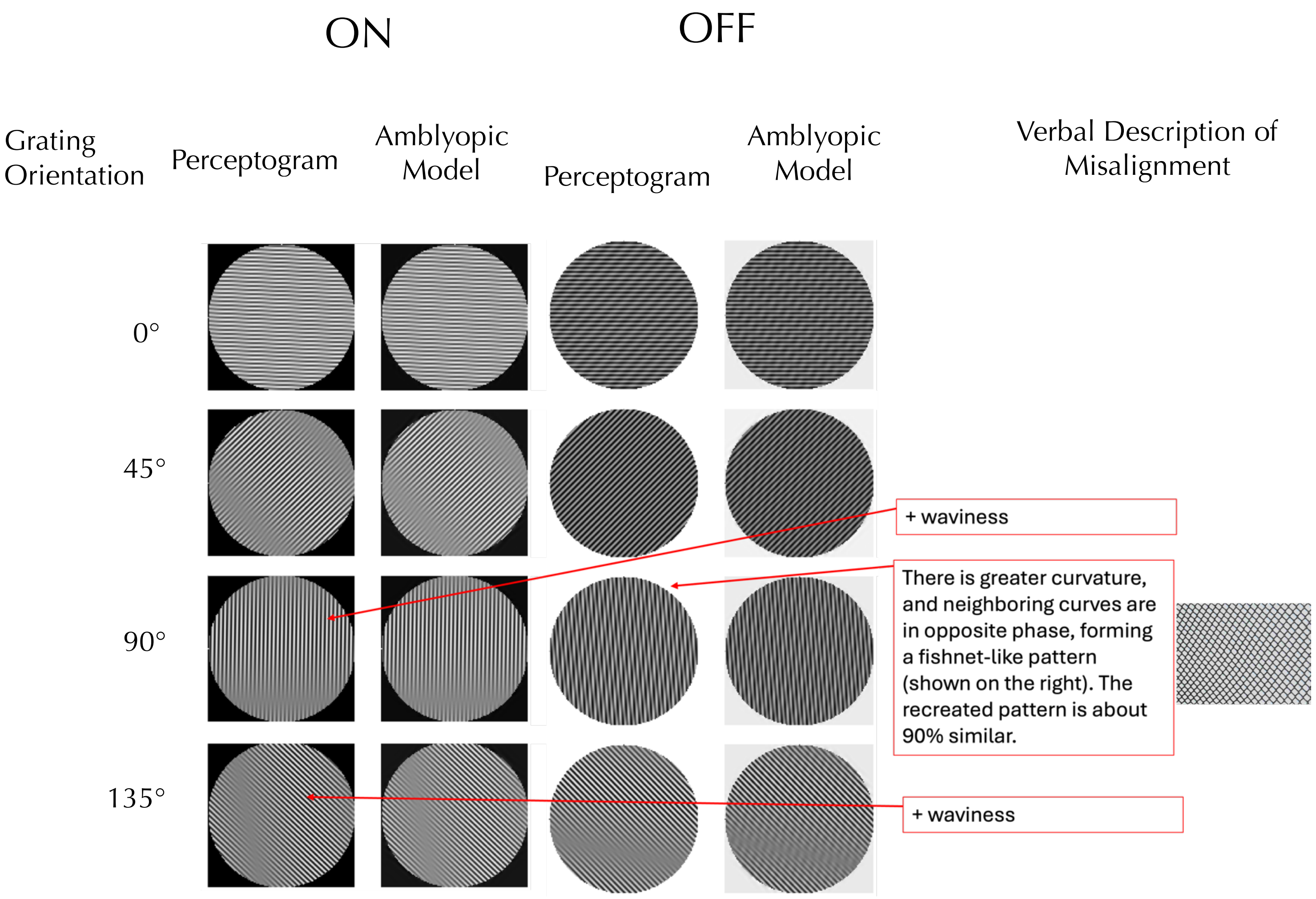

### 6 cycle/deg (Obs 5 Session 2)

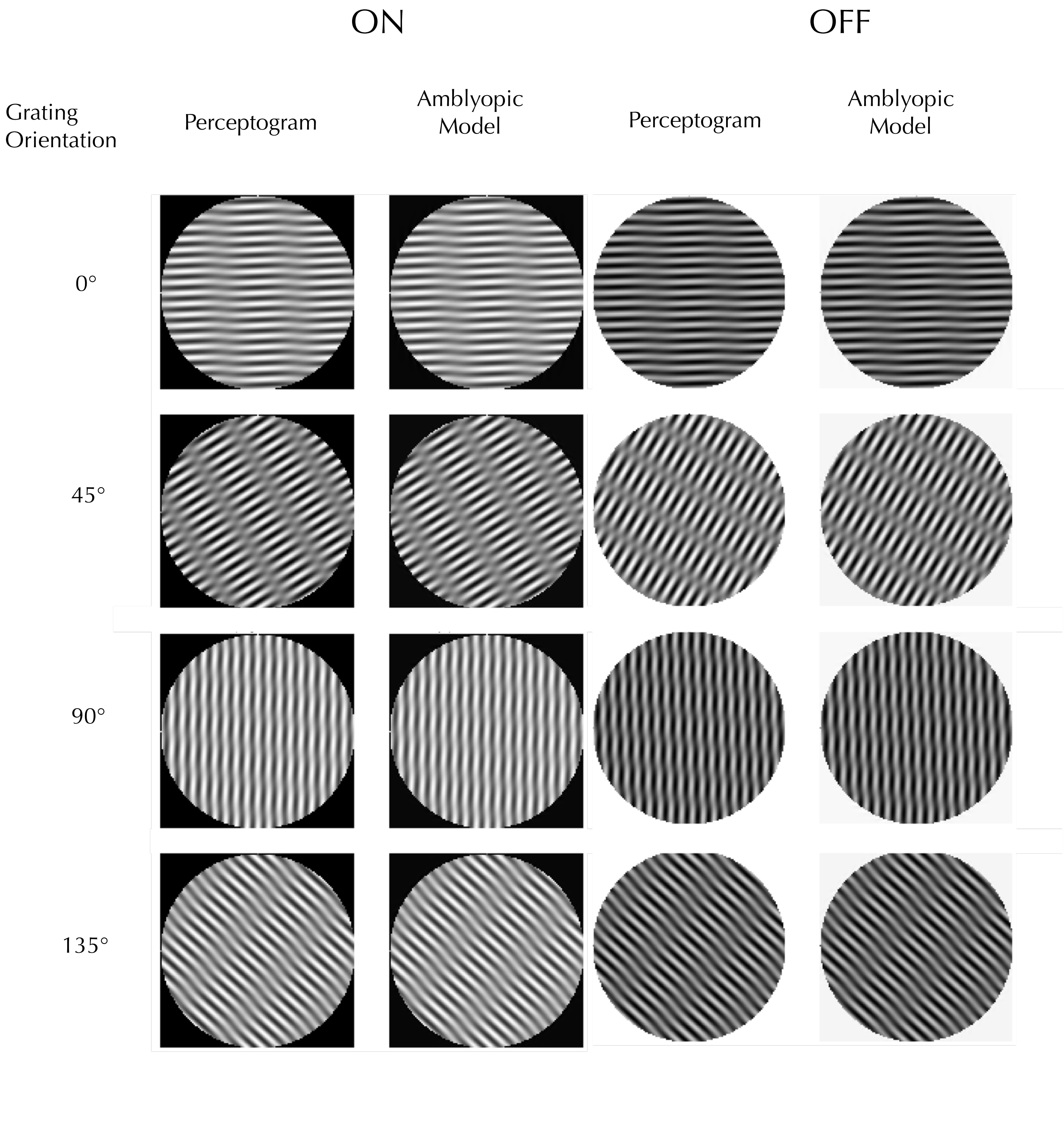

### 12 cycle/deg (Obs 5 Session 2)

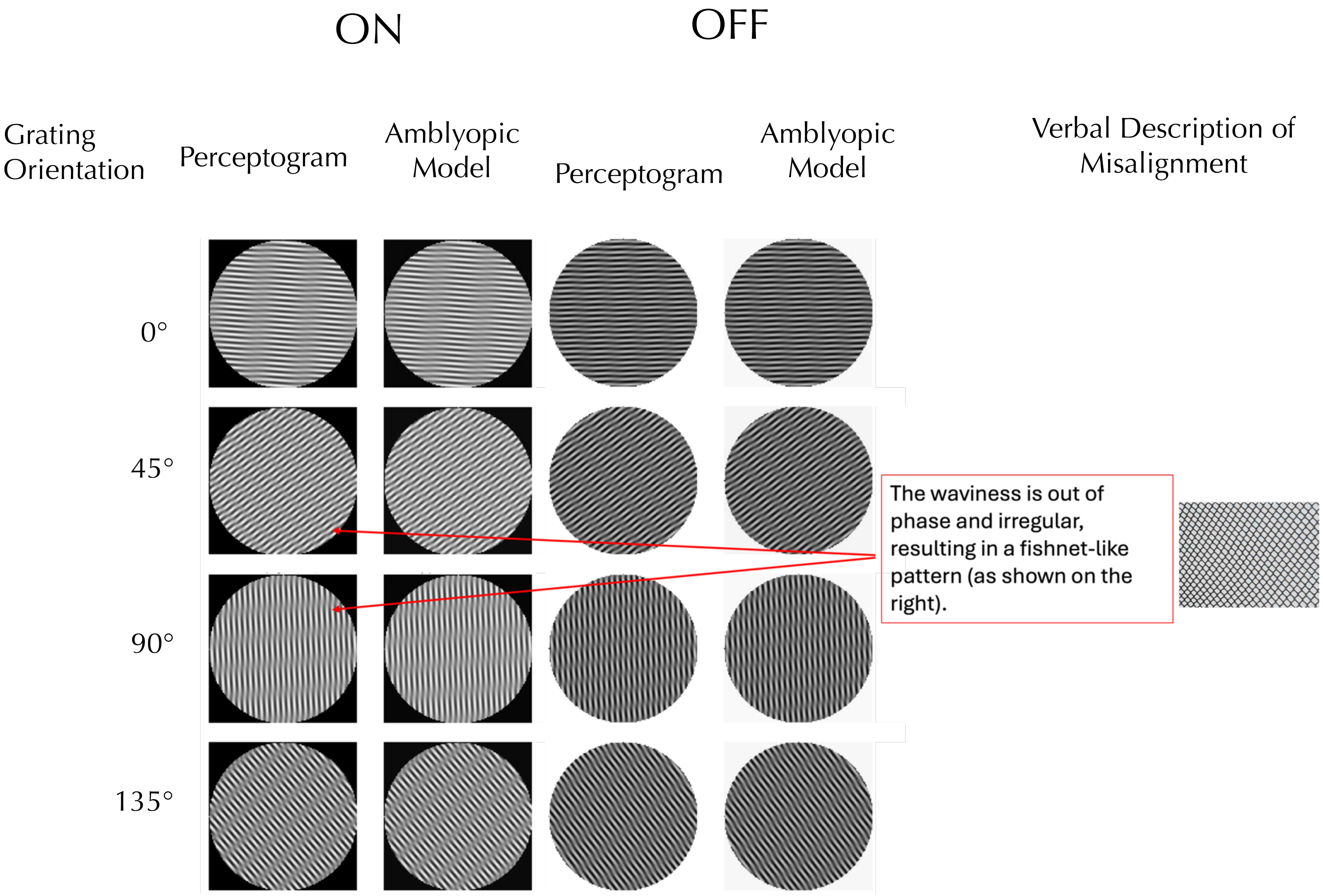

### 6 cycle/deg (Obs 5 Session 3)

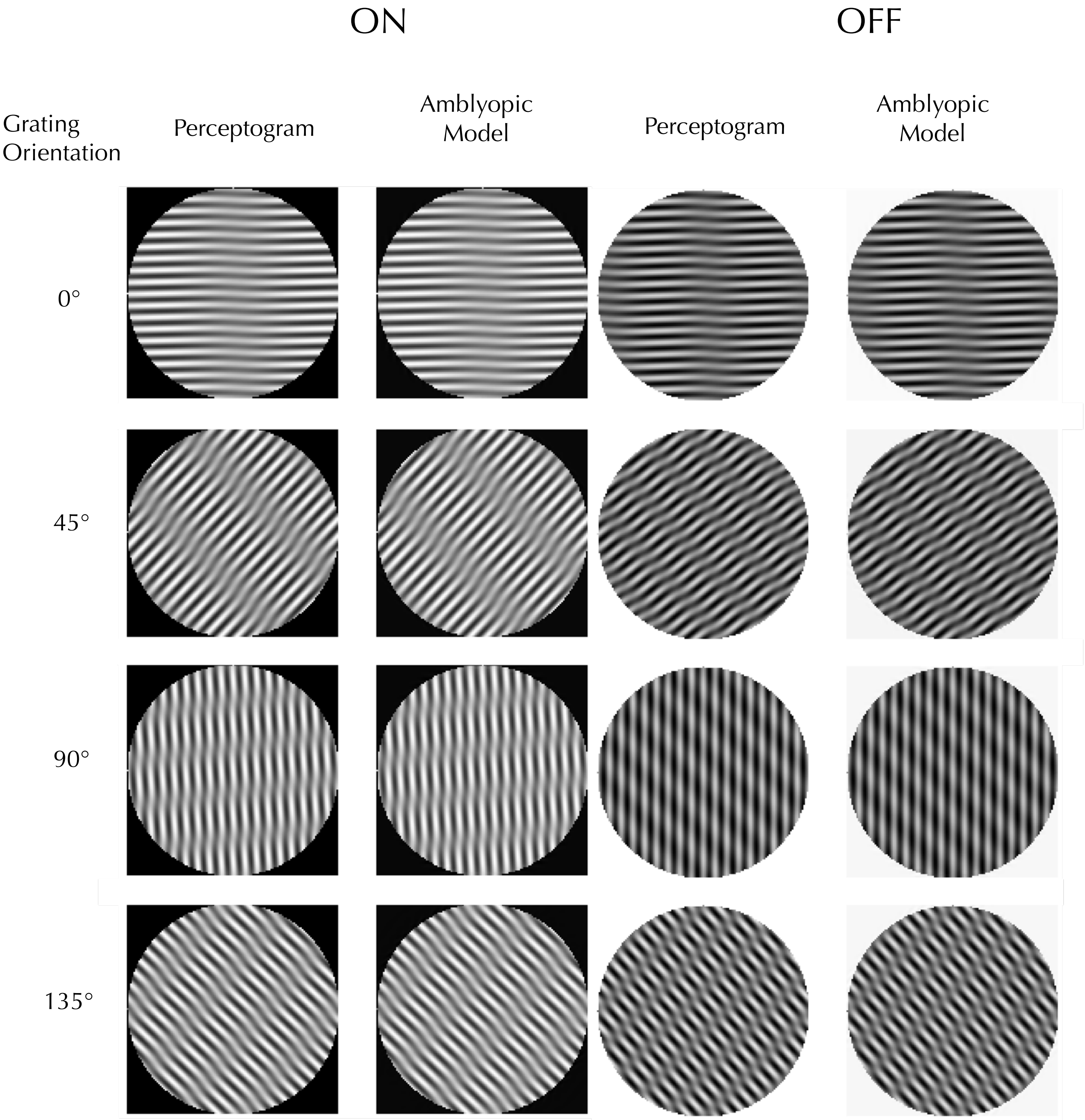

### 9 cycle/deg (Obs 5 Session 3)

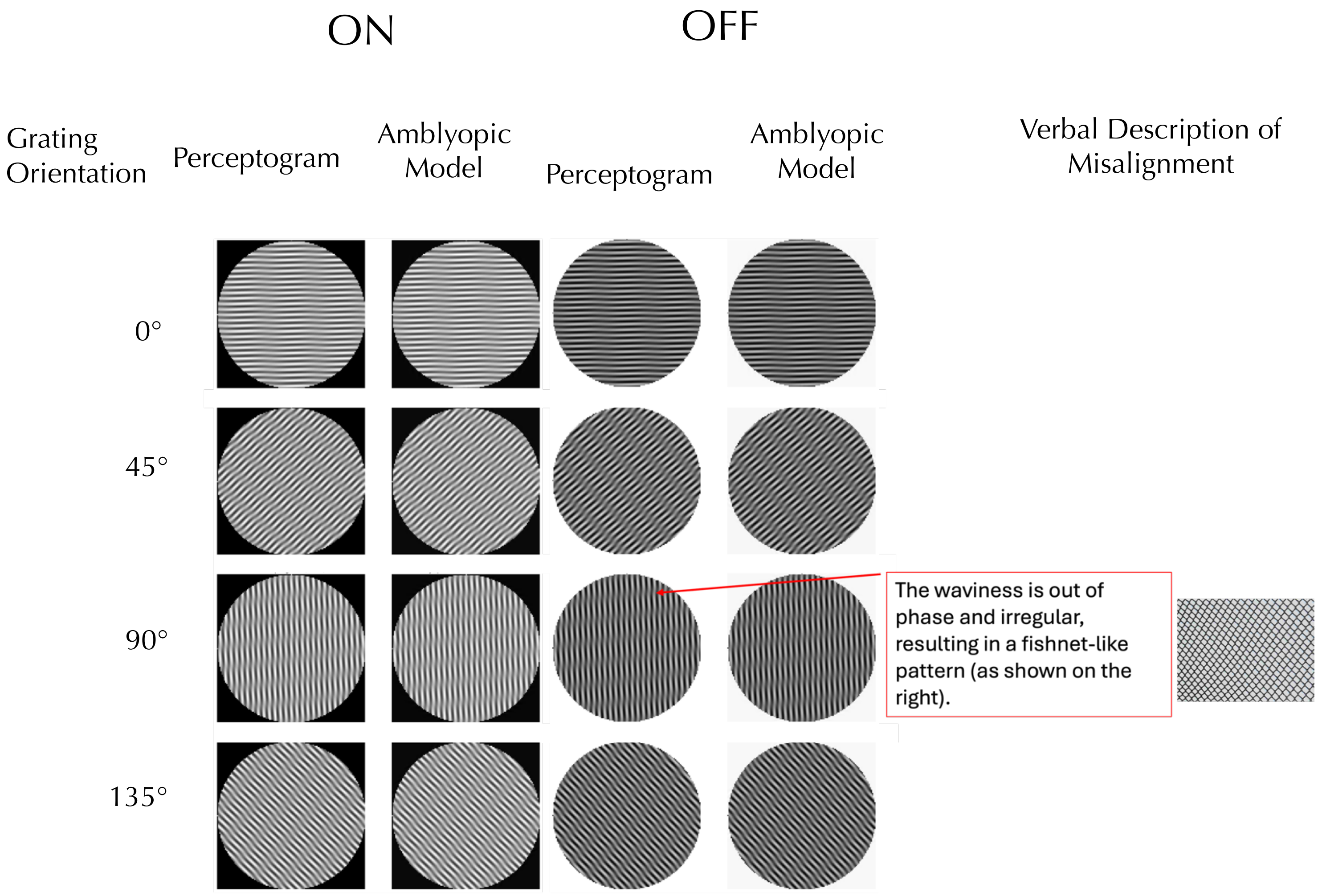

### 12 cycle/deg (Obs 5 Session 3)

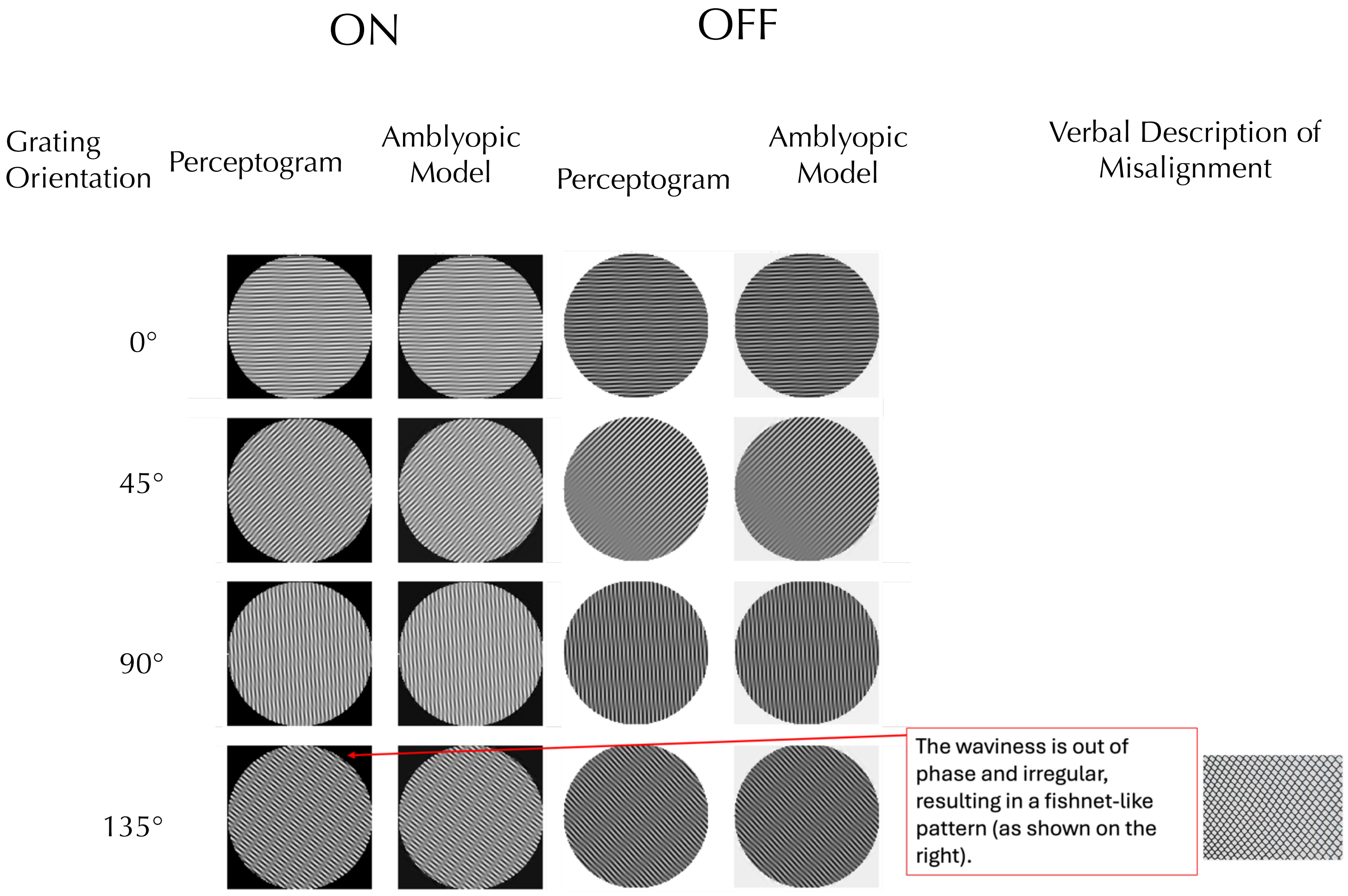


**Figure S1. Visual comparisons of the complete set of measured and generated Perceptograms:** In each panel, the left two columns display ON stimuli, and the right two OFF stimuli. Each row corresponds to a different stimulus orientation. Within each pair, the left image represents the observer's perceptual match, while the right image shows the corresponding reconstruction generated by the transformed AE cortex filters. In 200 out of 216 conditions, the generated plaids matched the perceived distortions. In the other 16 conditions, observers provided verbal descriptions of the mismatch. When verbal descriptions were insufficient, observers traced their percepts using Procreate on an iPad.
